# Transient induction of cell cycle promoter Fam64a improves cardiac function through regulating Klf15-dependent cardiomyocyte differentiation in mice

**DOI:** 10.1101/2021.04.01.438048

**Authors:** Ken Hashimoto, Aya Kodama, Momoko Ohira, Misaki Kimoto, Reiko Nakagawa, Yuu Usui, Yoshihiro Ujihara, Akira Hanashima, Satoshi Mohri

**Affiliations:** First Department of Physiology, Kawasaki Medical School, Kurashiki, 701-0192 Japan; Laboratory for Phyloinformatics, RIKEN Center for Biosystems Dynamics Research (BDR), Kobe, 650-0047, Japan; Department of Electrical and Mechanical Engineering, Nagoya Institute of Technology, Nagoya, 466-8555 Japan

**Keywords:** cardiomyocyte, regeneration, proliferation, differentiation, dedifferentiation, Fam64a, Klf15

## Abstract

The introduction of fetal or neonatal signatures such as cell cycle promoting genes into damaged adult hearts has been vigorously pursued as a promising strategy for stimulating proliferation and regeneration of adult cardiomyocytes, which normally cannot divide. However, cell division of cardiomyocytes requires preceding dedifferentiation with sarcomere disassembly and calcium dysregulation, which, in principle, compromises contractile function. To overcome this intrinsic dilemma, we explored the feasibility of optimizing the induction protocol of the cell cycle promoter in mice. As a model of this approach, we used Fam64a, a fetal-specific cardiomyocyte cell cycle promoter that we have recently identified. We first analyzed transgenic mice maintaining long-term cardiomyocyte-specific expression of Fam64a after birth, when endogenous expression was abolished. Despite having an enhanced proliferation of postnatal cardiomyocytes, these mice showed age-related cardiac dysfunction characterized by sustained cardiomyocyte dedifferentiation, which was reminiscent of the dilemma. Mechanistically, Fam64a inhibited glucocorticoid receptor-mediated transcriptional activation of Klf15, a key regulator that drives cardiomyocyte differentiation, thereby directing cardiomyocytes toward immature undifferentiated states. In contrast, transient induction of Fam64a in cryoinjured wildtype adult mice hearts improved functional recovery with augmented cell cycle activation of cardiomyocytes. These data indicate that optimizing the intensity and duration of the stimulant to avoid excessive cardiomyocyte dedifferentiation could pave the way toward developing efficient strategy for successful heart regeneration.

## Introduction

The limited proliferation potential of adult cardiomyocytes (CMs) is a major obstacle hindering regeneration of myocardium lost following injury. The introduction of fetal-specific signatures such as cell cycle promoting genes into damaged adult hearts has recently emerged as a promising strategy for stimulating CM proliferation (Borden et al., 2019; Mohamed et al., 2018; Nakada et al., 2017). This is because fetal CMs are highly proliferative and show a striking regenerative capacity following ablation of up to 60% of CMs (Sturzu et al., 2015). However, cell division of CMs requires preceding dedifferentiation that accompanies sarcomere disassembly and calcium dysregulation (D’Uva et al., 2015; Zhu et al 2020), which, in principle, compromises contractile function. Relatively little attention has been paid to this intrinsic dilemma, which has caused cardiac maladaptations in several settings that aimed at promoting CM proliferation. For example, persistent delivery of microRNA-199a in infarcted pig hearts resulted in uncontrolled cardiac repair characterized by CM dedifferentiation and proliferation, which led to sudden arrhythmic death (Gabisonia et al., 2019). Prolonged CM-specific inactivation of Hippo pathway during pressure overload led to cardiac dysfunction accompanied by CM dedifferentiation and proliferation in mice (Ikeda et al., 2019). Oncostatin M, which induces CM dedifferentiation and DNA synthesis, contributed to dilated cardiomyopathy when continuously activated (Kubin et al., 2011). Moreover, several unexplained results have been reported in which robust CM proliferation following injury did not contribute to heart regeneration (Marshall et al., 2019; Stockdale et al., 2018), potentially implying the expansion of immature non-functional CMs. Thus, understanding the mechanisms of these phenomena is crucial to overcome the intrinsic dilemma.

Here we tested the feasibility of optimizing the induction protocol of cell cycle promoting genes to find an effective way to circumvent the dilemma. As a model of this approach, we utilized Fam64a (family with sequence similarity 64, member A; also known as Pimreg, Cats, or Rcs1), a fetal-specific CM cell cycle promoter that we have recently identified (Hashimoto et al., 2017). The strong Fam64a expression in fetal CM nuclei was almost completely lost in postnatal CMs from mice (Hashimoto et al., 2017) and sheep (Locatelli et al., 2020). Fam64a knockdown inhibited and its overexpression enhanced fetal CM proliferation (Hashimoto et al., 2017). High expression of Fam64a has been noted in various types of tumors and this expression was well correlated with poor prognosis, suggesting an oncogenic potential of Fam64a (Jiang et al., 2019; Wei et al., 2019; Yao et al., 2019). Knockdown of Fam64a reduced proliferation in several cell lines (Barbutti et al., 2016; Jiang et al., 2019; Yao et al., 2019). These data led to the consistent view that Fam64a is a general cell cycle promoter. We first analyzed transgenic (TG) mice maintaining long-term CM-specific postnatal expression of Fam64a, when endogenous expression was abolished. These mice demonstrated cardiac dysfunction characterized by CM dedifferentiation, re-induction of fetal genes (known as a fetal gene program; FGP), and perturbation of the cardiac rhythm, despite an enhancement of CM proliferation, which was attributable to the intrinsic dilemma. The FGP is typically implicated in pathological cardiac remodeling that accompanies CM dedifferentiation and proliferation (Cui et al., 2018).

We explored the mechanisms underlying the FGP-mediated dedifferentiation and the rhythm disturbance induced in TG mice by focusing on Klf15 (Krüppel-like factor 15), a transcription factor that regulates diverse biological processes (Fan et al., 2018). Klf15 has been reported to promote differentiation (Asada et al., 2011; Mallipattu et al., 2012), and inhibit proliferation (Ray and Pollard, 2012; Yoda et al., 2015) in several cell types, including CMs (Zhang et al., 2015), suggesting the role in counteracting the FGP. Indeed, Klf15 inhibits pathological cardiac remodeling through repression of various transcription factors involved in the FGP (Fisch et al., 2007; Leenders et al., 2010, 2012). Loss of Klf15 relieves this inhibitory effect, leading to heart failure (Leenders et al., 2010). Klf15 is also a principal regulator that establishes cardiac rhythmicity (Jeyaraj et al., 2012a; Zhang et al., 2015). A deficiency or excess of Klf15 perturbs rhythmic CM electrical activity and increases susceptibility to ventricular arrhythmias (Jeyaraj et al., 2012a). It also controls other rhythmic biological processes, such as bile acid synthesis (Han et al., 2015) and nitrogen homeostasis (Jeyaraj et al., 2012b).

In the present study, we show that Fam64a inhibits Klf15 activity by glucocorticoid receptor (GR)-mediated transcriptional regulation, which results in sustained FGP-driven dedifferentiation coupled with rhythm disturbance in TG mice. Thus, we propose a novel function of Fam64a in directing CMs toward immature undifferentiated states by inhibiting Klf15, which, in the long-term, could lead to cardiac dysfunction. In contrast, transient induction of Fam64a in the cryoinjured hearts of wildtype (WT) adult mice showed progressively improved functional recovery with minimum left ventricular dilation and augmented CM cell cycle activation. These data indicate that optimizing the induction protocol, e.g., the intensity and duration, of the cell cycle stimulant, particularly to avoid excessive CM dedifferentiation, is a promising strategy to overcome the intrinsic dilemma, and holds great promise to attain cardiac regeneration.

## Results

### Enhanced CM proliferation at the neonatal, adult, and aged stages in CM-specific Fam64a TG mice

We have established CM-specific Fam64a TG mice under the control of the alpha myosin heavy chain promoter with a C-terminal FLAG tag (Figure 1–figure supplement 1A–C, Figure 1–source data 1). Expressed protein was confirmed to localize in the CM nuclei, in the same location as an endogenous protein (Hashimoto et al., 2017) (Figure 1–figure supplement 1D). We found that TG mice showed greater proliferative capacity than WT mice at the neonatal (postnatal day, P12–P15), adult (6–7 wks), and aged (> 25 wks) stages, as indicated by increased positivity for Ki67, a cell cycle marker, and phospho-histone H3 (pH3), a mitosis marker (Figure 1A–C). Cell cycle promoting genes were consistently upregulated in TG mice (Figure 1D–F). These in vivo results are in agreement with our previous in vitro analyses identifying Fam64a as a CM cell cycle promoter (Hashimoto et al., 2017).

**Figure 1.**
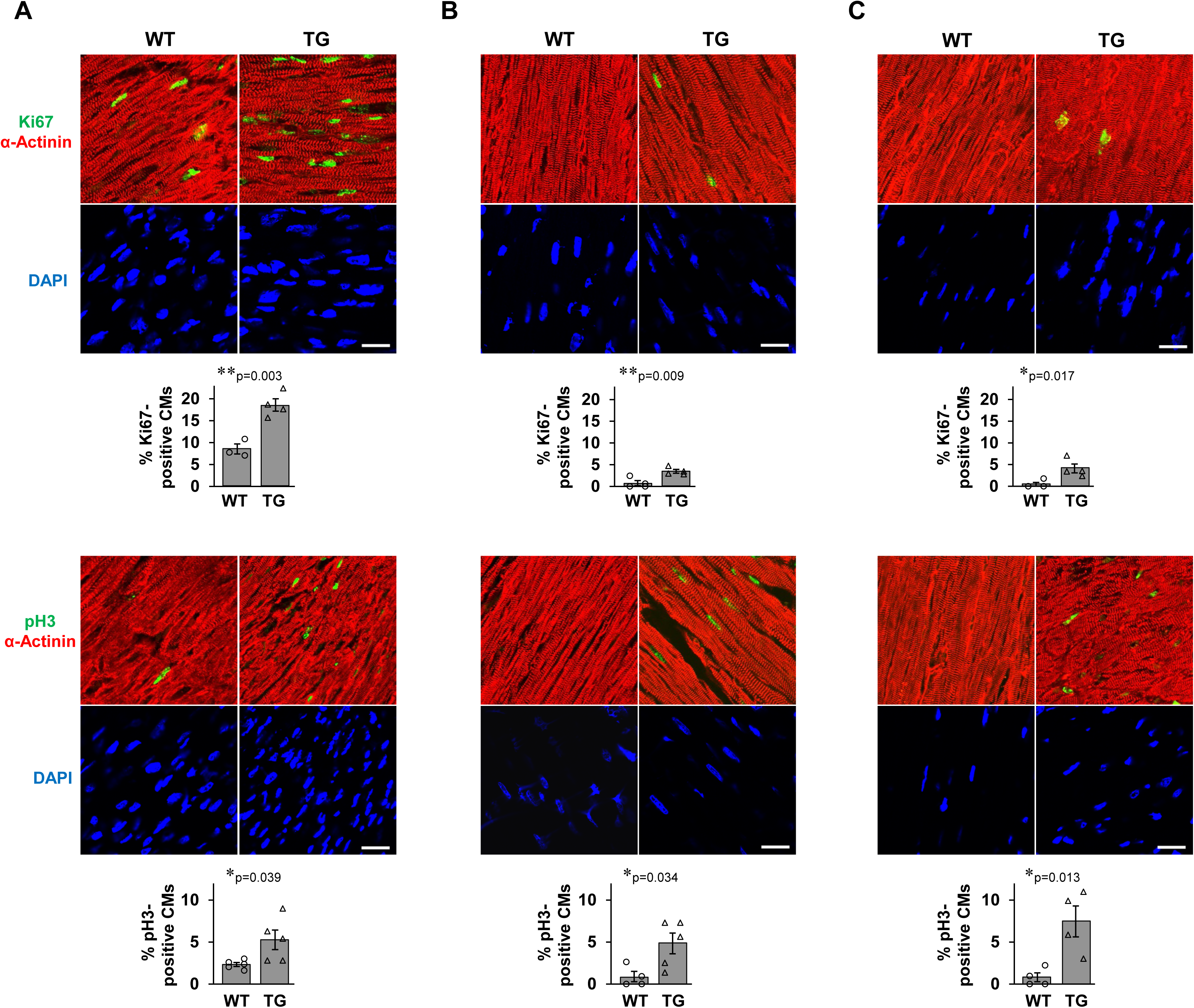
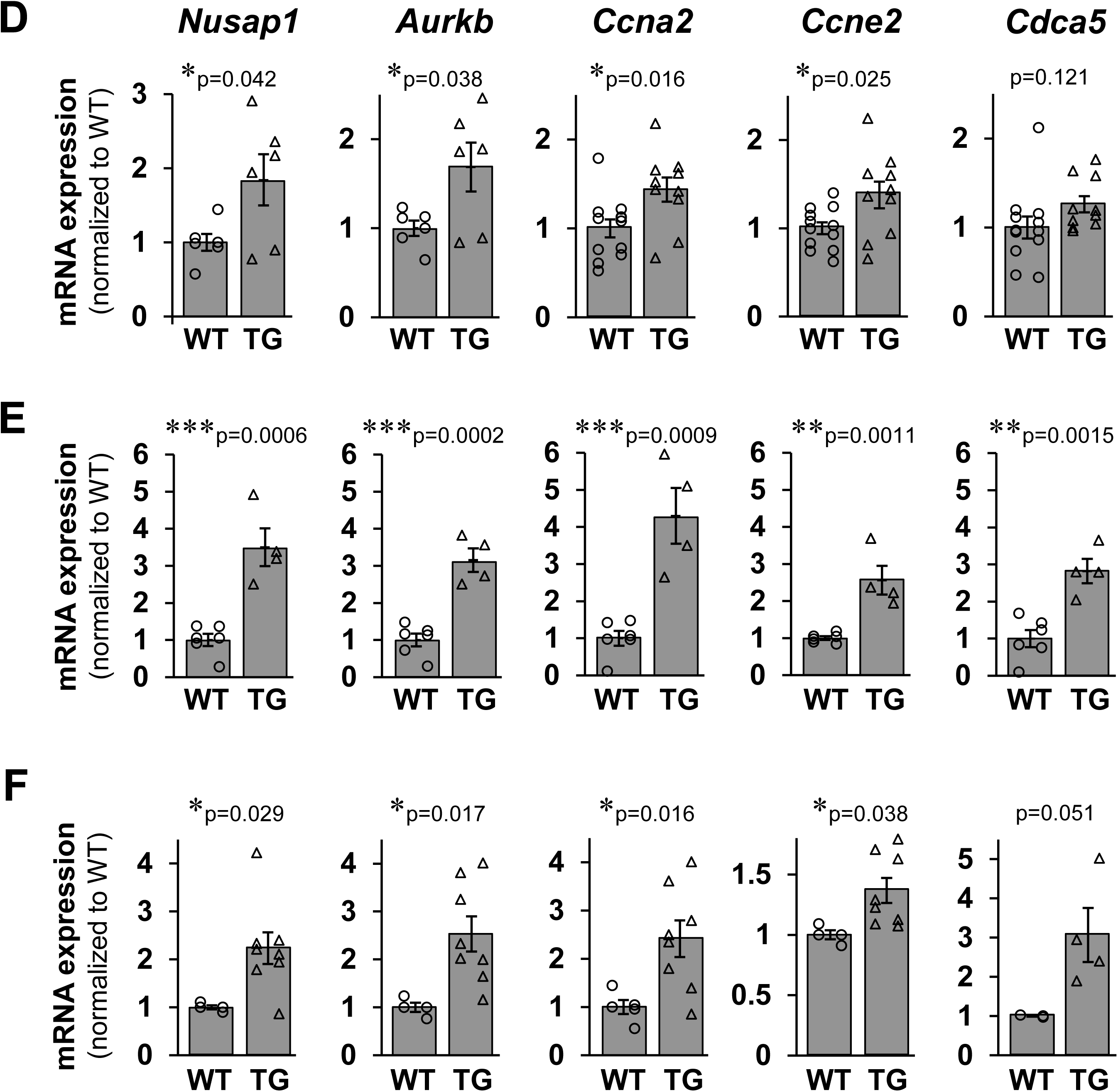
Enhanced CM proliferation at the neonatal, adult, and aged stages in CM-specific Fam64a TG mice. **A–C.** Immunofluorescence for Ki67 (upper panel, green) and phospho-histone H3 (pH3) (lower panel, green) observed in sarcomeric α-actinin (red, as a CM marker) and DAPI (blue) in WT and TG mice heart sections at neonatal (**A,** P12–P15), adult (**B**, 6–7 wks), and aged (**C**, > 25 wks) stages. Quantitative analysis for the percentage of Ki67-positive and pH3-positive CMs were shown on the bottom of each image. n = 3–5 mice per group. For each mouse, 400–800, 80–160, and 80–230 CMs were counted for neonatal, adult, and aged stages, respectively. * p < 0.05 and ** p < 0.01 as compared to WT by Student’s two-tailed unpaired t-test. Error bar = SEM. Scale bar = 20 µm. **D–F.** qPCR analysis of cell cycle promoting genes in WT and TG mice hearts at neonatal (**D,** P12–P15), adult (**E**, 6–7 wks), and aged (**F**, > 25 wks) stages. Data were shown as normalized to WT. n = 3–12 mice per group. * p < 0.05, ** p < 0.01, *** p < 0.001 as compared to WT by Student’s two-tailed unpaired t-test. Error bar = SEM.

### Fam64a TG mice show age-related cardiac dysfunction with poor survival

Echocardiography demonstrated that TG mice showed progressive left ventricular dilation both at diastole and systole, leading to a severe decline in cardiac contractile function as estimated by fractional shortening when compared to WT mice (Figure 2A–D). Histological assessment indicated that although there was no apparent difference at the neonatal stage, chamber dilation and wall thinning in left ventricle was progressively observed in TG mice in the adult and aged stages (Figure 2E–G). Survival analysis demonstrated a marked drop in survival rate in TG mice (Figure 2H). These data show that Fam64a TG mice develop age-related cardiac dysfunction with poor survival despite their enhanced CM proliferation.

**Figure 2.**
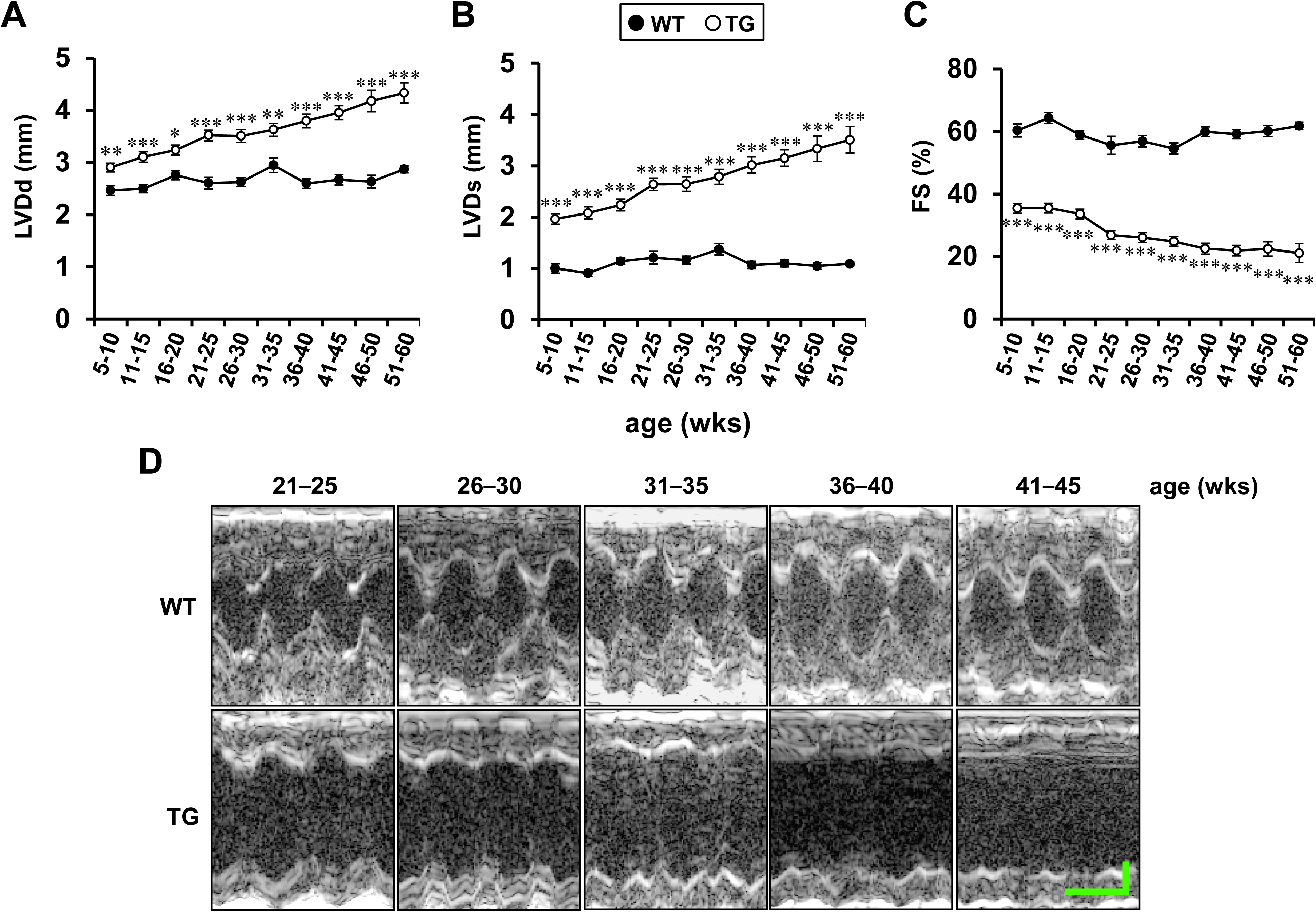
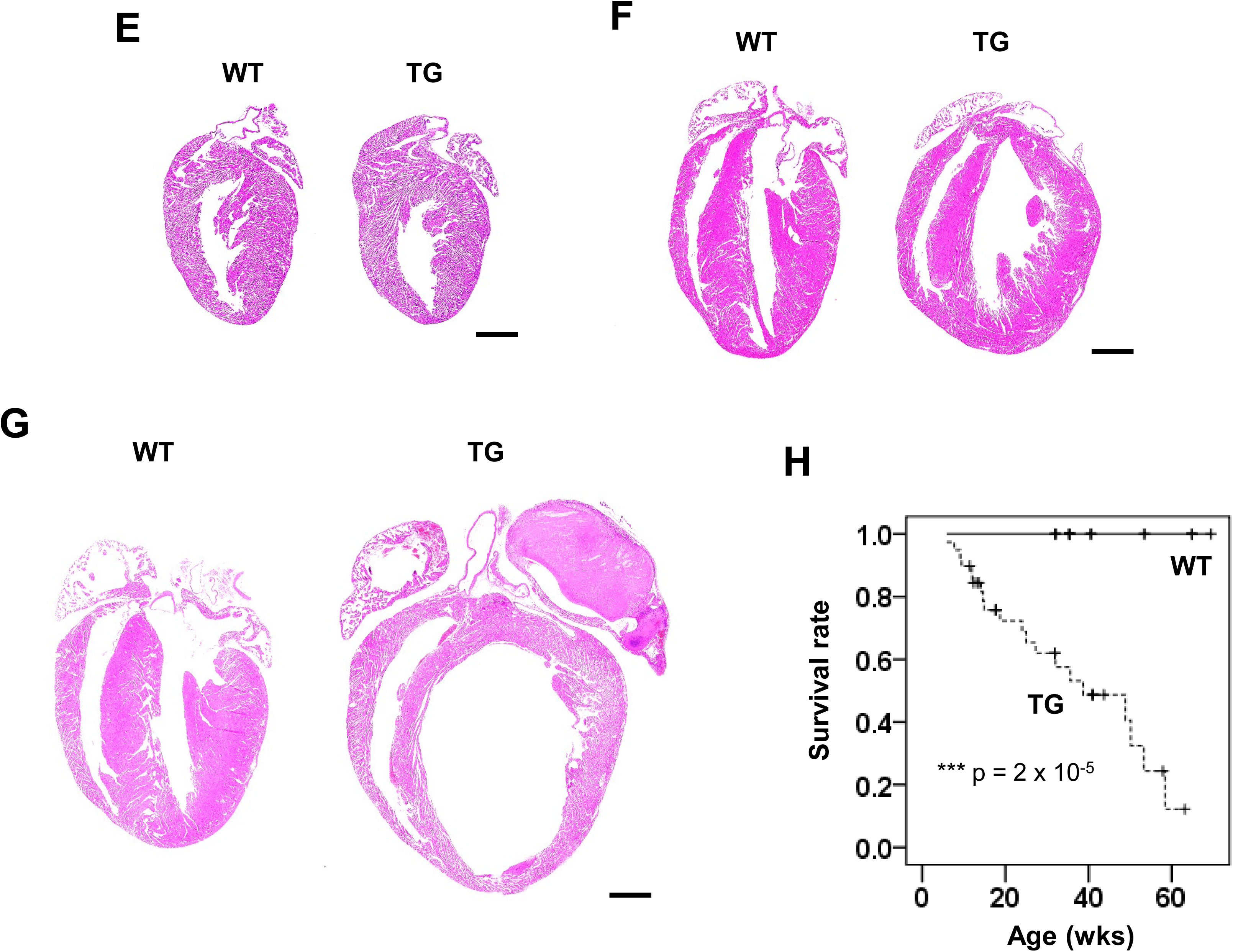
Fam64a TG mice show age-related cardiac dysfunction with poor survival. **A–D.** Left ventricular internal diameter at end diastole (LVDd, **A**) and end systole (LVDs, **B**) were measured in sedated WT (filled circle) and TG (open circle) mice from 5 to 60 wks of age by two-dimensional transthoracic M-mode echocardiography. Fractional shortening (FS, **C**) was calculated as ([LVDd–LVDs]/LVDd)×100 (%), and was used as an index of cardiac contractile function. Representative tracings were shown in **D** (Horizontal scale bar = 100 ms, vertical scale bar = 1 mm). In WT mice, no significant change was observed in LVDd, LVDs, and FS over the course of the study, with the only exception of LVDd at 31–35wks significantly larger as compared to 5–10 wks (One-way ANOVA with Tukey’s post hoc test). By contrast in TG mice, LVDd and LVDs were significantly increased, and FS was significantly decreased at 21–25 wks and afterwards as compared to 5–10 wks (One-way ANOVA with Tukey’s post hoc test). Consequently, in TG mice, LVDd and LVDs were significantly larger and FS was significantly smaller at all stage as compared to WT mice of the same age (* p < 0.05, ** p < 0.01, and *** p < 0.001 as compared to WT by Student’s two-tailed unpaired t-test.). n = 17–83 mice per group at each age which partially includes repetitive measurements of the same animal at different age. Error bar = SEM. **E–G.** Representative H&E staining for WT and TG mice heart sections at neonatal (**E,** P12–P15), adult (**F**, 6–7 wks), and aged (**G**, > 25 wks) stages. Scale bar = 1 mm. **H.** Overall survival curves of WT and TG mice were analyzed by Kaplan–Meier method. The vertical line in each plot indicates the censored data. n = 21–39 mice per group. *** p < 0.001 as compared to WT by log-rank test.

### Cardiac dysfunction in Fam64a TG mice is attributable to impaired Ca^2+^ handling and contractile property in CMs

Calcium homeostasis in CMs is essential for the maintenance of normal cardiac function. The Ca^2+^ transient measurements conducted in isolated CMs from aged mice (29–32 wks) revealed a reduction in the peak amplitude and a delay in the time to peak, indicating impaired Ca^2+^ mobilization in TG mice as compared to WT mice (Figure 3A–C). Although not statistically significant, a tendency was observed toward an increased time constant during the decay phase (Figure 3D) and a decreased sarcoplasmic reticulum (SR) Ca^2+^ content (Figure 3E), suggesting impaired Ca^2+^ re-uptake into the SR in TG mice. Cell shortening in response to electrical stimuli was decreased in TG mice at all the frequencies tested, indicating impairment of the CM contractile properties (Figure 3F). These findings were corroborated by qPCR analysis showing that many of the principal genes involved in Ca^2+^ handling in mature differentiated CMs, including *Ryr2*, *Cacna1c*, *Atp2a2*, and *Thra*, were downregulated in TG mice at the neonatal, adult, and aged stages (Figure 3G–I). Similar downregulations were observed in primary cultures of isolated fetal CMs overexpressing Fam64a (Figure 3–figure supplement 1). These data suggest that impaired Ca^2+^ handling and CM contractile properties with a reduction of mature cell markers, which can be regarded as a sign of CM dedifferentiation, contribute to cardiac dysfunction in Fam64a TG mice.

**Figure 3.**
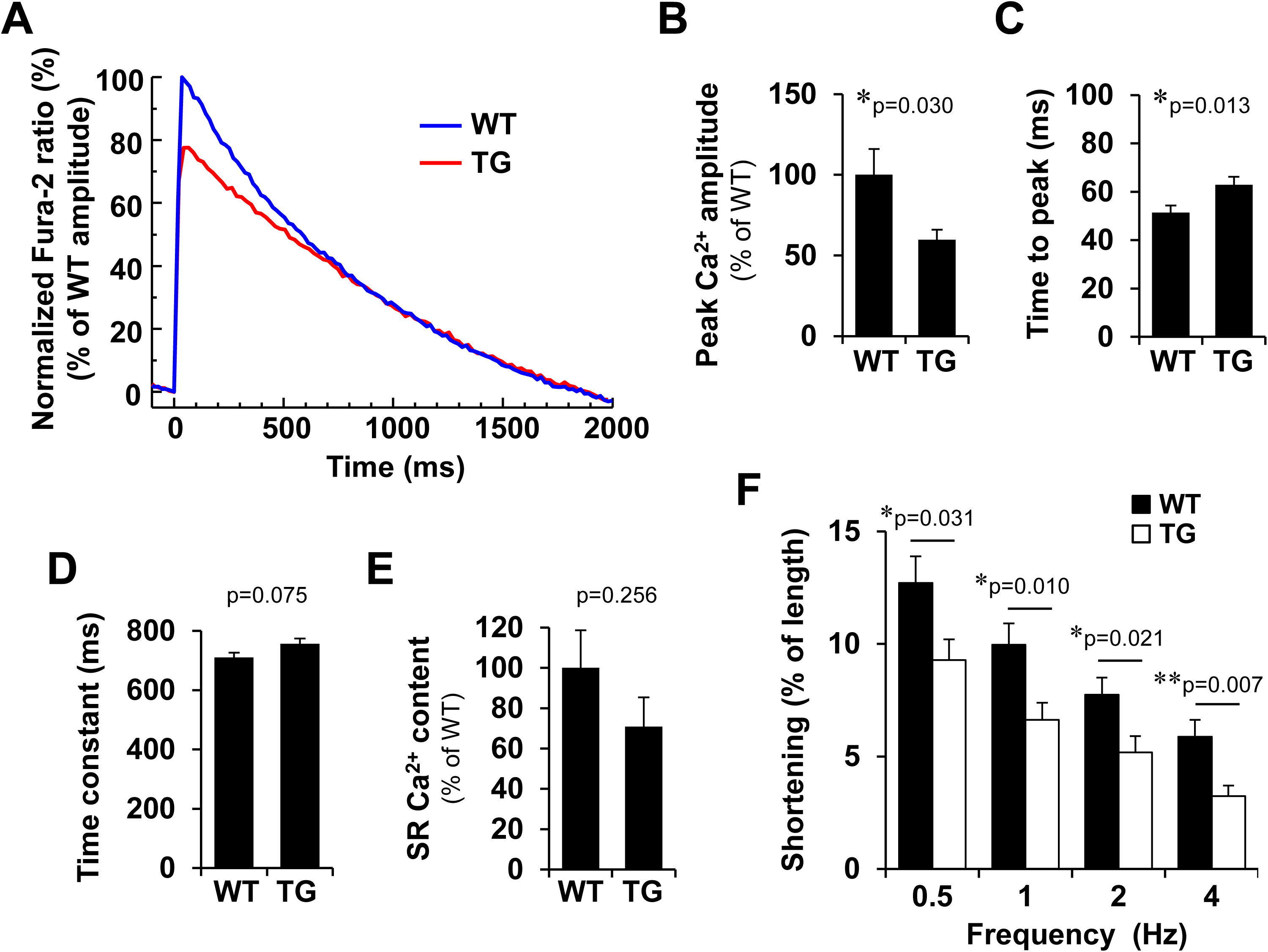
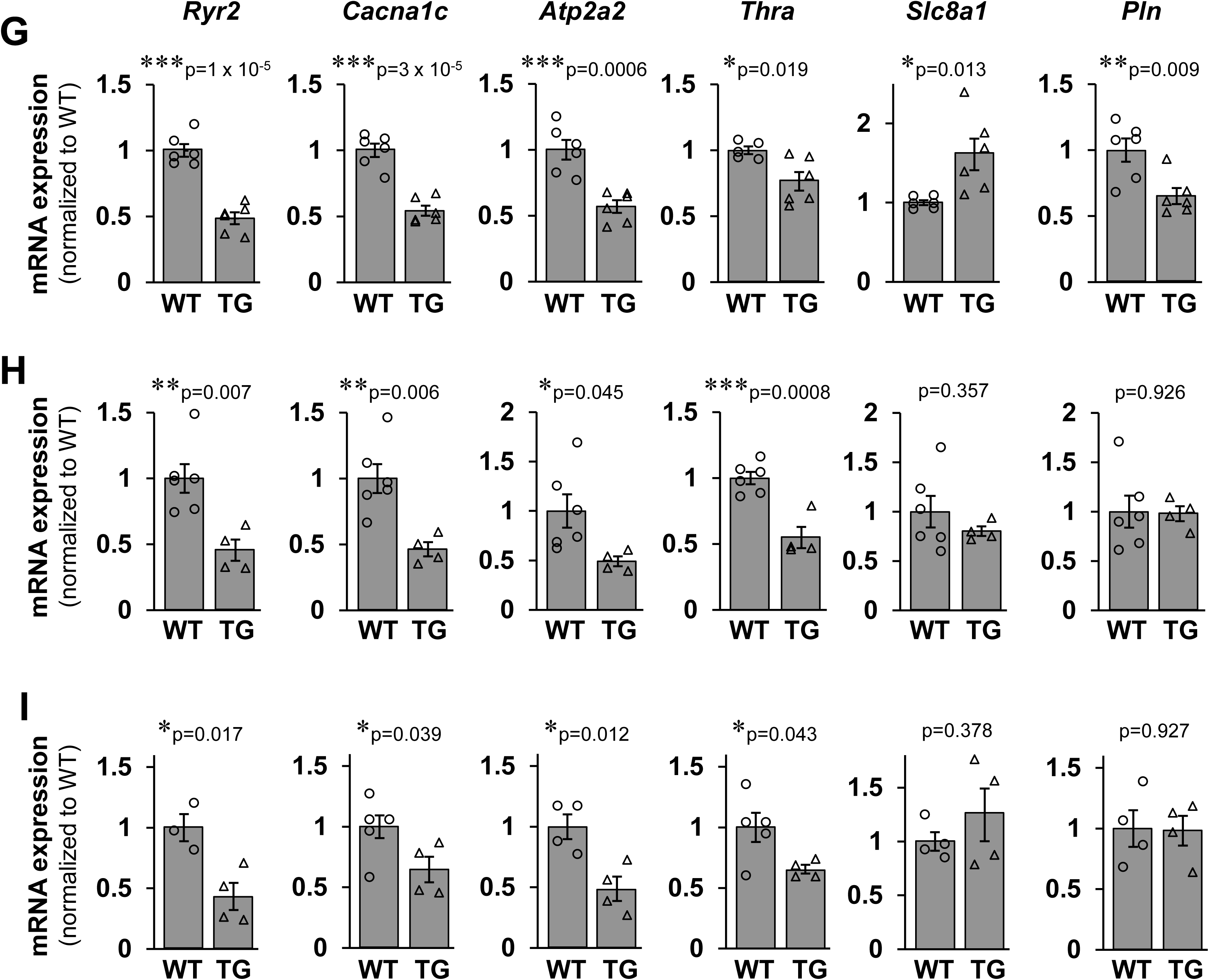
Cardiac dysfunction in Fam64a TG mice is attributable to impaired Ca^2+^ handling and contractile property in CMs. **A–F**. Ca^2+^ transients and cell shortening were measured in isolated CMs from WT and TG mice at aged stages (29–32 wks). Representative Fura-2 ratio tracings of CMs (WT: blue, TG: red) stimulated at 0.5 Hz were shown as normalized to the peak value in WT set at 100 % (**A**). Quantitative analysis for peak Ca^2+^ amplitude (% normalized to WT) (**B**), time to peak (**C**), time constant (**D**), and sarcoplasmic reticulum (SR) Ca^2+^ content (% normalized to WT) (**E**) were shown. Cell shortening (% of initial cell length) stimulated at indicated frequencies were shown in (**F**). WT: filled bar, TG: open bar. n = 9–28 CMs from 3 WT mice and 23–50 CMs from 2 TG mice. In **B–E**, * p < 0.05 as compared to WT by Student’s two-tailed unpaired t-test. In **F**, * p < 0.05 and ** p < 0.01 as compared to WT under the same stimulating frequency by Student’s two-tailed unpaired t-test. Error bar = SEM. **G–I.** qPCR analysis of Ca^2+^ handling genes in WT and TG mice hearts at neonatal (**G,** P12–P15), adult (**H**, 6–7 wks), and aged (**I**, > 25 wks) stages. Data were shown as normalized to WT. n = 3–6 mice per group. * p < 0.05, ** p < 0.01, *** p < 0.001 as compared to WT by Student’s two-tailed unpaired t-test. Error bar = SEM.

### Fam64a TG mice show signs of CM dedifferentiation and the FGP

In contrast to the highly organized sarcomere structures with small, rod-shaped nuclei observed in WT mice, the TG mice displayed disorganization of the sarcomere structures with enlarged, irregular-shaped nuclei frequently observed, indicative of CM dedifferentiation (D’Uva et al., 2015; Ikeda et al., 2019; Jopling et al., 2010; Kubin et al., 2011) (Figure 4A). We also found a decreased CM cell size in TG mice, another sign of CM dedifferentiation, as estimated by two-dimensional image analysis of freshly isolated CMs (Figure 4B). The qPCR analysis demonstrated a strong induction of a suite of genes involved in the FGP and dedifferentiation in TG mice, including *Myh7*, *Nppa*, *Nppb*, *Tnnt1*, *Acta2*, *Myl4*, *Cacna1h*, and *Pi16* (Chiang et al.,2009; Cui et al., 2018; Ikeda et al., 2019; Kubin et al., 2011; Peng et al., 2017; Taegtmeyer et al., 2010) (Figure 4C). Conversely, mRNA level was decreased for connexin 43 (*Gja1*), a primary component of the mature gap junction (Figure 4C). Moreover, genes encoding several K^+^ channel subunits were consistently repressed, which is recognized as a sign of CM dedifferentiation (Karbassi et al., 2020) (Figure 4–figure supplement 1). The reduction of thyroid hormone receptor α (*Thra*) (Figure 3G–I) was also considered a sign of dedifferentiation because thyroid hormone T3 is a strong inducer of postnatal CM differentiation (Karbassi et al., 2020). These data indicate that CM dedifferentiation and the FGP are induced in Fam64a TG mice.

**Figure 4.**
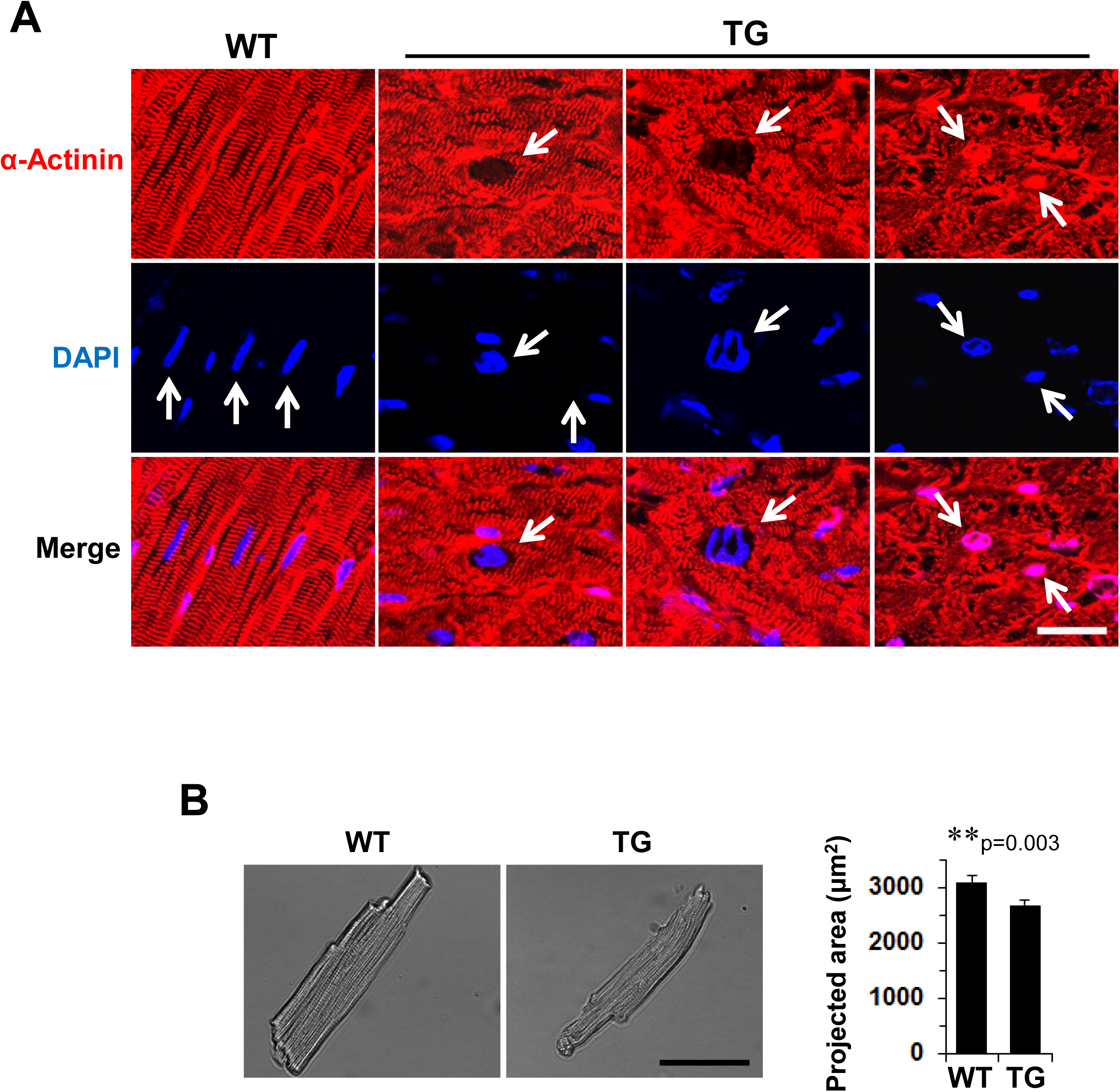
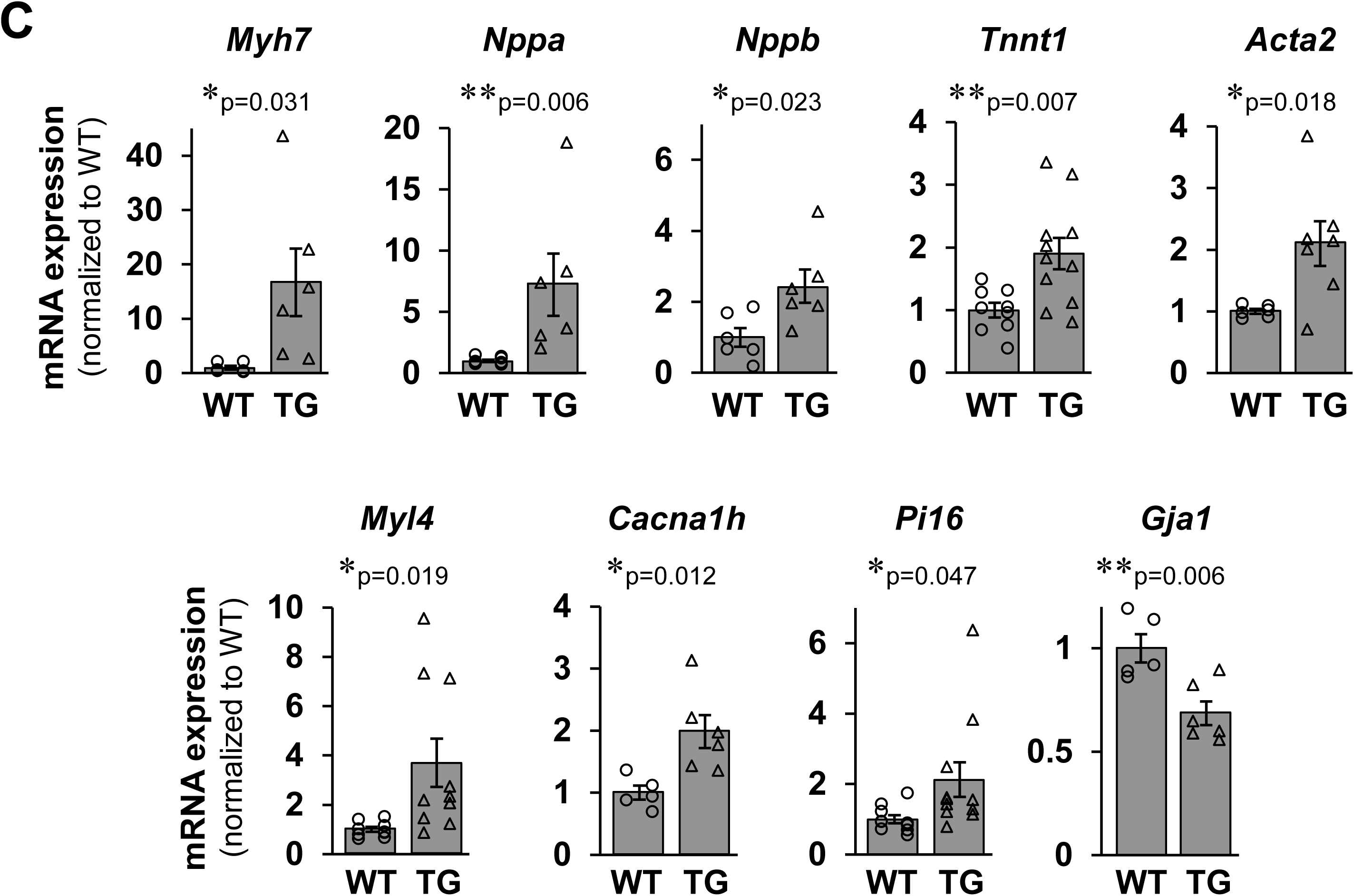
Fam64a TG mice show signs of CM dedifferentiation and the FGP. **A.** Immunofluorescence for sarcomeric α-actinin (red) and DAPI (blue) in WT and TG mice heart sections at > 25 wks aged stages. In WT mice, highly organized sarcomere structure with small, rod-shaped nuclei (arrows) was observed. In contrast, disorganization of sarcomeres with enlarged, irregular-shaped nuclei were frequently observed in TG mice (arrows). Scale bar = 20 µm. **B.** Representative image of freshly isolated CMs from WT and TG mice at > 25 wks aged stages, obtained by differential interference contrast optics. CM cell size was evaluated as a two-dimensional projected area. n = 75 CMs from 3 WT mice and 113 CMs from 2 TG mice. ** p < 0.01 as compared to WT by Student’s two-tailed unpaired t-test. Error bar = SEM. Scale bar = 50 µm. **C.** qPCR analysis of genes involved in CM dedifferentiation and the FGP, and a mature gap junction component connexin 43 (*Gja1*) in WT and TG mice hearts. Data were shown as normalized to WT. n = 5–11 mice per group. * p < 0.05 and ** p < 0.01 as compared to WT by Student’s two-tailed unpaired t-test. Error bar = SEM.

### Perturbed cardiac rhythmicity and locomotor activity in Fam64a TG mice

Surprisingly, genome-wide RNA-seq analysis of heart samples identified a pathway called circadian rhythm as the most differentially regulated pathway in TG mice, as indicated by an enrichment score far greater than other pathways like cardiac muscle contraction, hypertrophic cardiomyopathy, and dilated cardiomyopathy (Figure 5A; RNA-seq data have been deposited in DDBJ sequencing read archive (DRA) under the accession number DRA009818, http://trace.ddbj.nig.ac.jp/DRASearch). Some of the principal genes relating to circadian rhythm, including *Arntl* (known as Bmal1), *Cry1*, *Per2*, *Npas2*, and *Dbp*, were dysregulated in TG hearts (Figure 5B). Telemetric measurements using freely moving conscious mice revealed perturbed heart rate regulation in the TG mice: (1) Heart rate was consistently low in TG mice, irrespective of daytime or nighttime, throughout the course of measurements of up to 8 days (Figure 5C). (2) Nighttime-to-daytime ratios of heart rate in TG mice were slightly, but significantly, lower than in WT mice, and had values of less than 1, indicating an abnormal daytime (inactive phase)-dominant heart rate regulation (Figure 5D). In addition, TG mice frequently developed premature ventricular contraction, either as a single form or more hazardous serial forms, in sharp contrast to WT mice that displayed virtually no such arrhythmias (Figure 5E, F). Decreased expression of connexin 43 could partially explain these aberrant phenotypes (Figure 4C).

**Figure 5.**
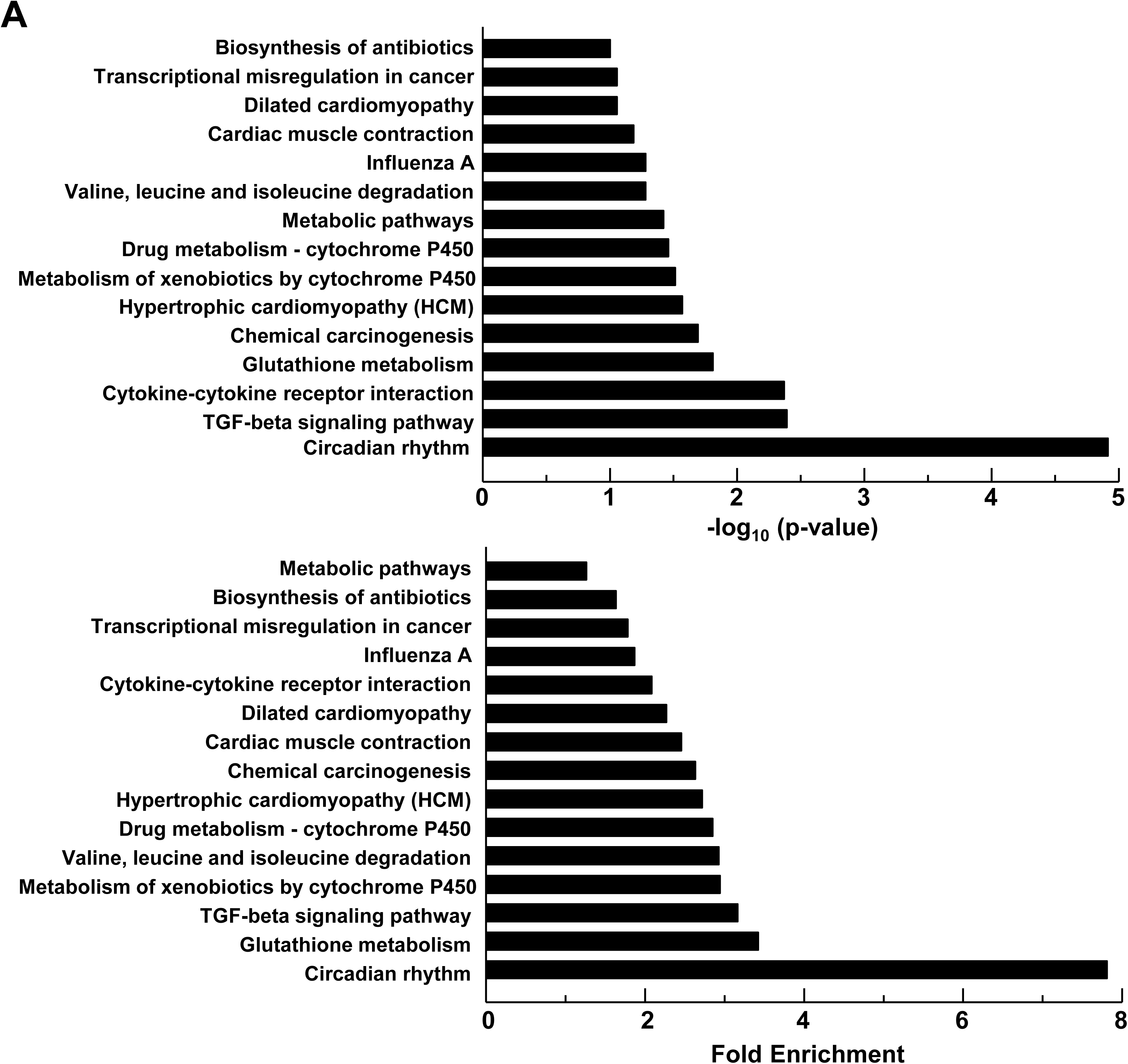
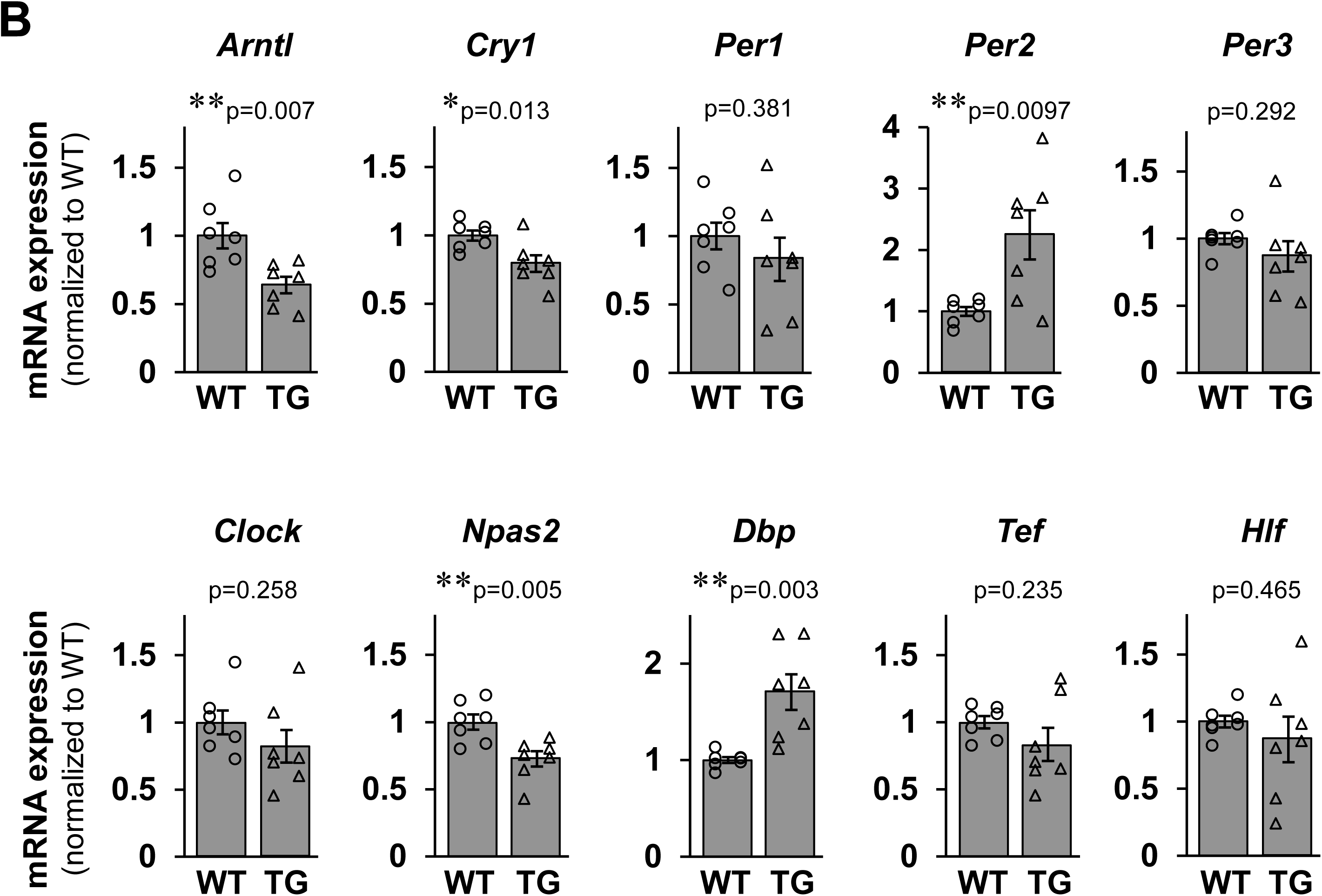
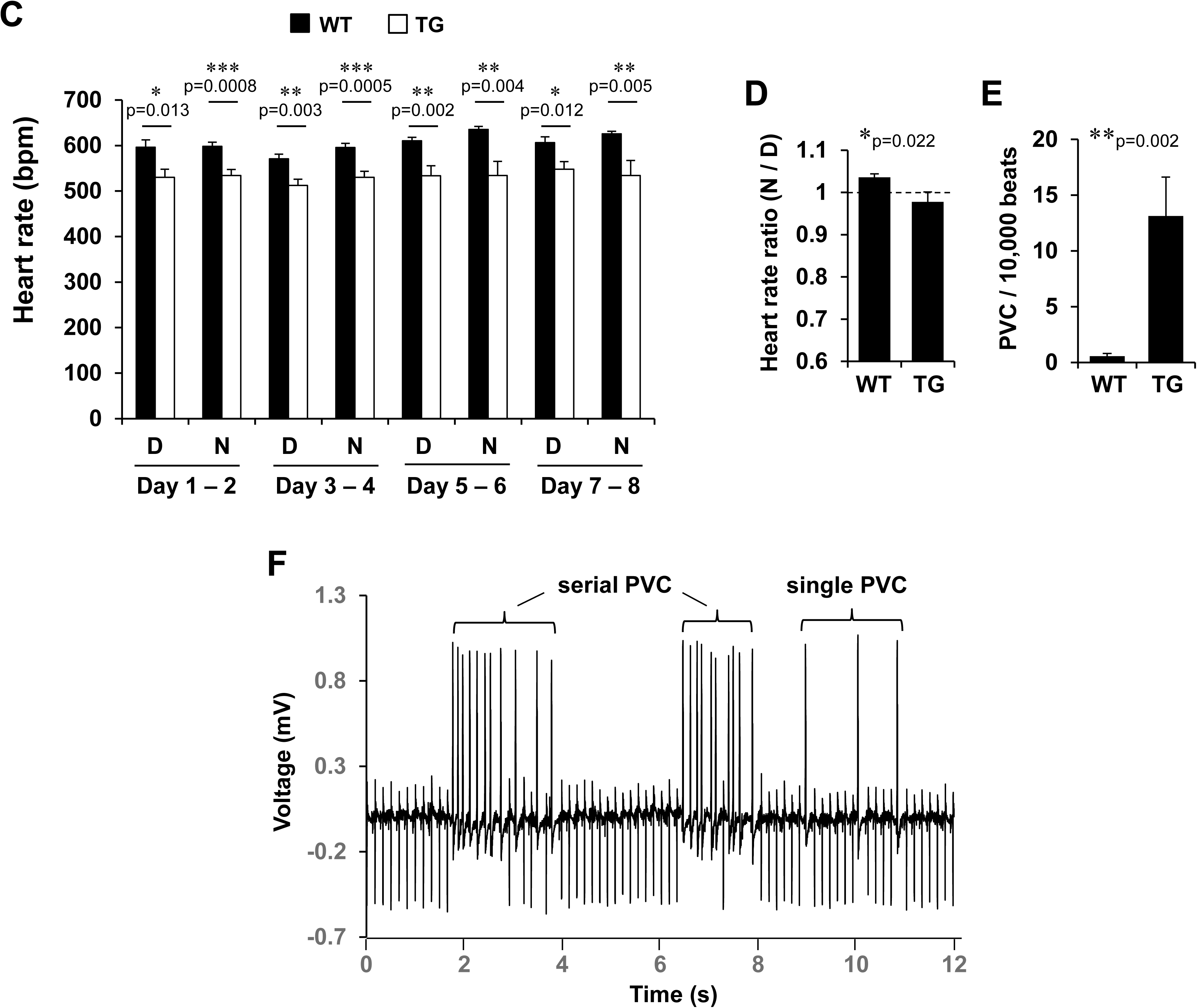
Rhythmic CM electrical activity is perturbed in Fam64a TG mice. **A.** Based on RNA-seq data comparing gene expressions in WT vs. TG mice hearts, functional annotation analysis was performed using DAVID. In this analysis, genes upregulating > 2.0 and downregulating < 0.5 in TG relative to WT mice were used to identify the differentially regulated gene pathways. The rank order of potency for p value (top) and fold enrichment score (bottom) were shown. **B.** qPCR analysis of genes involved in circadian rhythm in WT and TG mice hearts. Data were shown as normalized to WT. n = 7 mice per group. * p < 0.05, ** p < 0.01 as compared to WT by Student’s two-tailed unpaired t-test. Error bar = SEM. **C–F.** Telemetric ECG measurements using freely moving conscious mice for a total of 8 days in a 12-h light:12-h dark cycle (lights-on at 8 am). **C:** Averaged heart rate during daytime (D; 8 am to 8 pm) and nighttime (N; 8 pm to 8 am) in WT (filled bar) and TG (open bar) mice. Data pooled for every 2 days were shown. **D:** Nighttime (8 pm to 8 am)-to-daytime (8 am to 8 pm) ratio of heart rate in WT and TG mice. Data pooled for 8 days were shown. **E:** Frequency of premature ventricular contraction (PVC) per 10,000 beats in WT and TG mice. Data were analyzed for two representative time periods per animal. See methods for details. **F:** Representative ECG tracings in TG mice developing single and serial forms of PVC. Data were analyzed from 6 WT mice and 7 TG mice at adult and aged stages. * p < 0.05, ** p < 0.01, and *** p < 0.001 as compared to WT by Student’s two-tailed unpaired t-test. Error bar = SEM.

Locomotor activity analysis of mice using the infrared motion detector also demonstrated perturbation of rhythmic behavior in TG mice: Whereas the nighttime (active phase)-dominant activity was observed in both WT and TG mice, the activity in TG mice was decreased during nighttime and increased during daytime when compared to WT mice (Figure 5–figure supplement 1). The abnormal daytime-dominant heart rate regulation (Figure 5D) might be responsible for this phenotype. These data indicate that rhythmic CM electrical activity and locomotor activity were perturbed in Fam64a TG mice.

### Fam64a inhibits Klf15 activity by glucocorticoid receptor-mediated transcriptional regulation, thereby inducing the FGP-mediated CM dedifferentiation and the rhythm disturbance in TG mice

We explored the mechanisms underlying the FGP-mediated dedifferentiation and the rhythm disturbance induced in TG mice by focusing on Klf15, a key regulator that drives CM differentiation by repressing the FGP and establishing CM rhythmic activity (Fisch et al., 2007; Jeyaraj et al., 2012a; Leenders et al., 2010, 2012; Zhang et al., 2015). We found that mRNA expression of Klf15 and its downstream target Kcnip2 (Kv channel-interacting protein 2, also known as KChIP2) (Jeyaraj et al., 2012a) was strongly upregulated during the course of differentiation in WT hearts but was severely depressed in Fam64a-overexpressing TG mice hearts, suggesting that Fam64a inhibits Klf15 activity at the transcriptional level (Figure 6A). We conducted a comprehensive search for interacting partners of Fam64a that could mediate the inhibition of Klf15 by immunoprecipitation (Figure 6–figure supplement 1, Figure 6–source data 1), followed by mass spectrometry (Mass spectrometry data have been deposited in ProteomeXchange Consortium via jPOST, with the dataset identifiers PXD020570 and JPST000921. Preview code for reviewers https://repository.jpostdb.org/preview/497638005f1cd5ace27c1, Access key: 6340). This led us to focus on glucocorticoid receptor (GR), which was confirmed to form a complex with Fam64a in CMs (Figure 6B, Figure 6–source data 2). GR has previously been shown to bind to the promoter of Klf15 and stimulate its expression (Asada et al., 2011; Lee et al., 2016; Sasse et al., 2013).

**Figure 6.**
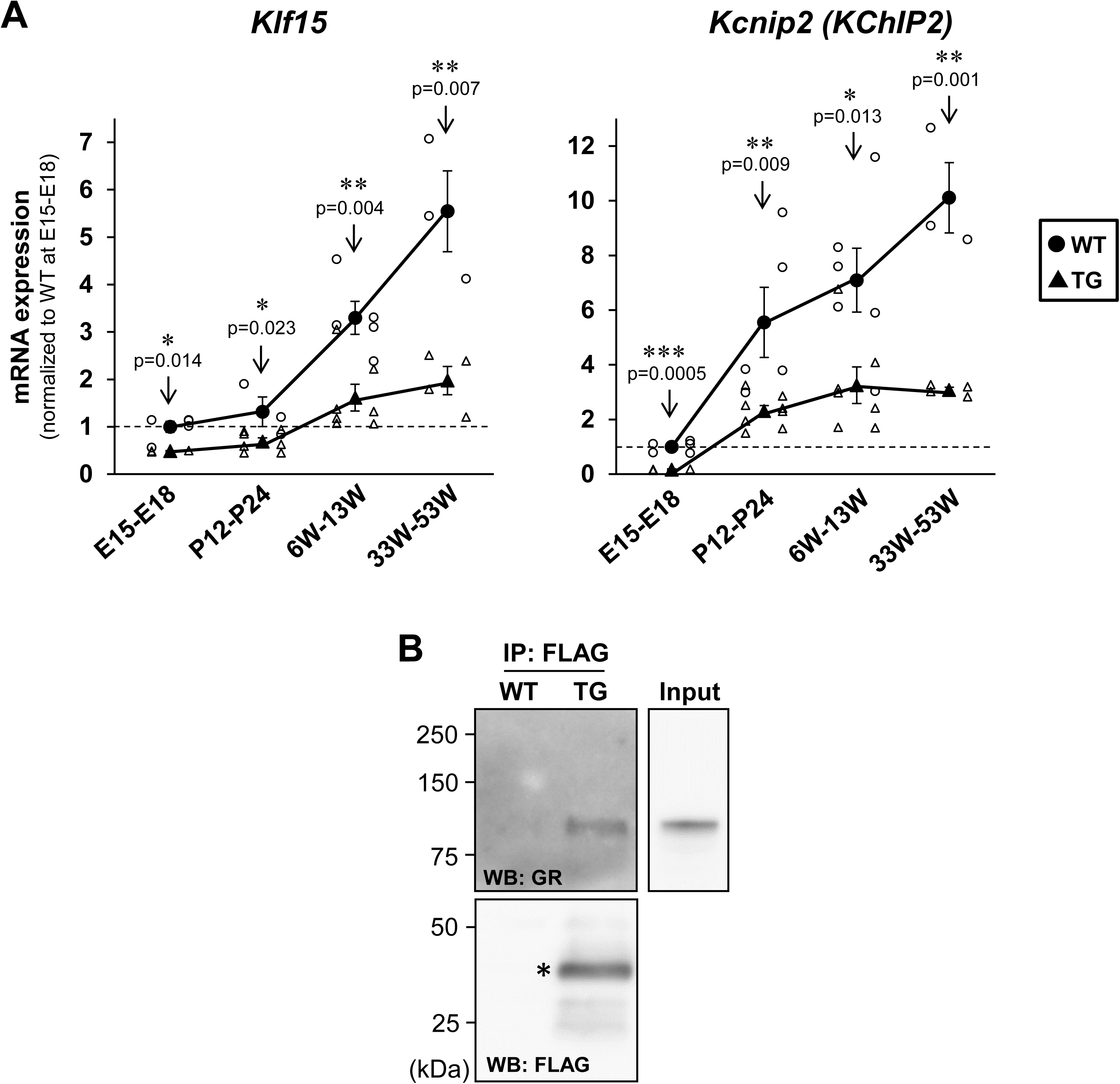
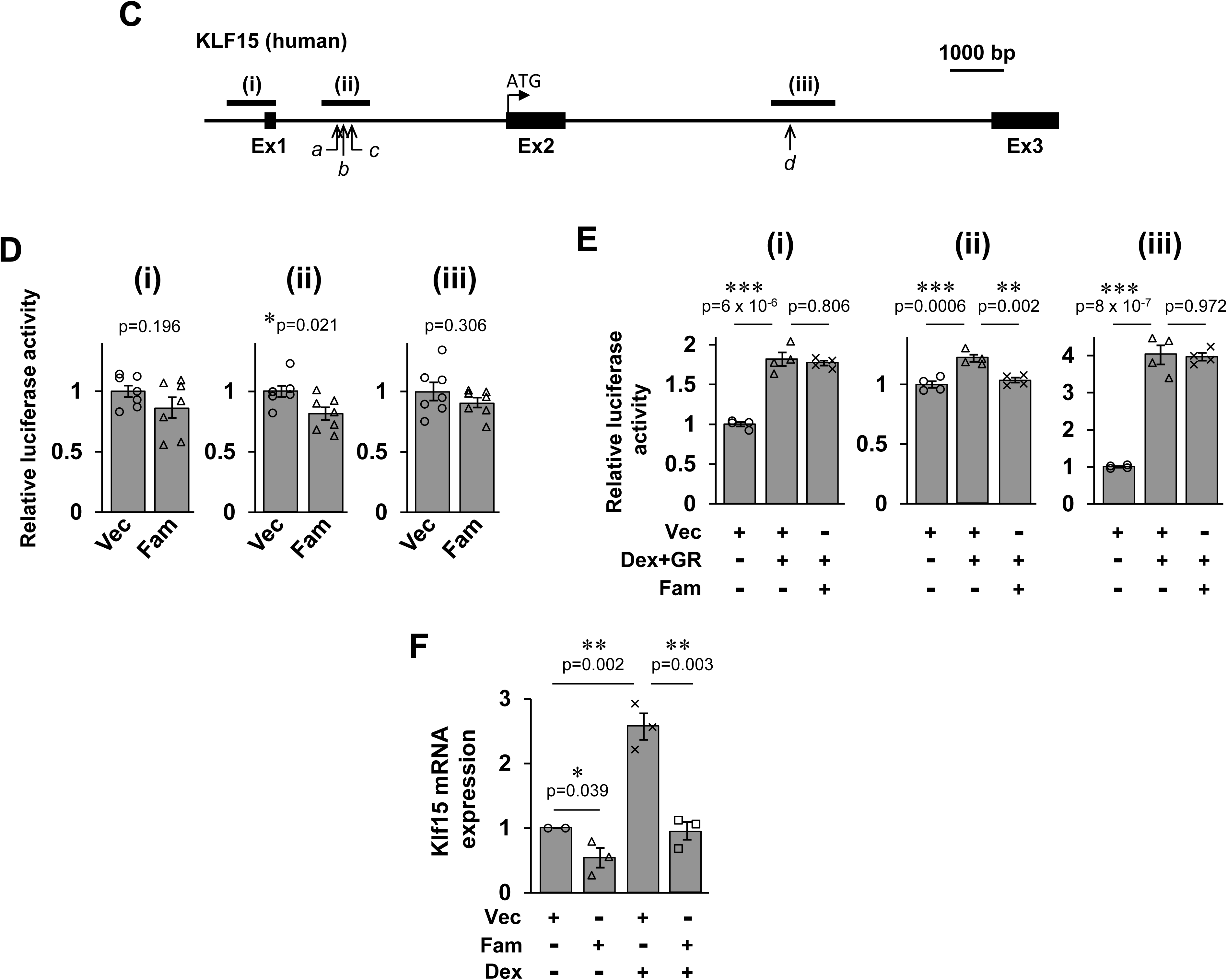
Fam64a inhibits Klf15 activity by glucocorticoid receptor-mediated transcriptional regulation, thereby inducing the FGP-mediated CM dedifferentiation and the rhythm disturbance in TG mice. **A.** qPCR analysis of *Klf15* and *Kcnip2* (KChIP2) at fetal, neonatal, adult, and aged stages from WT (circle) and TG (triangle) mice hearts. Data were shown as normalized to WT at fetal stage set at 1. In WT mice, Klf15 expression was significantly increased at 6W–13W and afterwards as compared to fetal stage (E15–E18) (One-way ANOVA with Tukey’s post hoc test). Likewise, Kcnip2 expression was significantly increased at P12–P24 and afterwards as compared to fetal stage (E15– E18) (One-way ANOVA with Tukey’s post hoc test). By contrast in TG mice, the expressions of both genes were significantly attenuated at all stage as compared to WT mice of the same age (* p < 0.05, ** p < 0.01, and *** p < 0.001 as compared to WT by Student’s two-tailed unpaired t-test.). n = 3–8 mice per group. Error bar = SEM. **B.** Immunoprecipitation (IP) against FLAG peptide that was expressed as a C-terminal tag of overexpressing Fam64a protein in TG mice hearts, followed by western blotting (WB) using glucocorticoid receptor (GR) antibody, which detected GR protein in TG, but not in WT mice heart lysates. This indicates that Fam64a forms complex with GR in CMs. Western blotting using FLAG antibody correctly detected Fam64a-FLAG fusion protein (*) in TG, but not in WT mice heart lysates, validating the immunoprecipitation procedure. **C.** Three reporter constructs on human KLF15 locus was used in luciferase reporter assay. The construct (ⅰ) contains common promoter sequence upstream of the first exon. The construct (ⅱ) and (ⅲ) contains three (marked as *a–c*) and the remaining one (marked as *d*), respectively, of the four GR binding sites previously reported. Ex = exon. **D.** HEK293T/17 cells were transiently transfected with Fam64a expression vector (Fam) or control empty vector (Vec), and one of the three reporter constructs (ⅰ–ⅲ). The luciferase activity of each reporter construct was normalized to that of the control reporter construct, and was expressed as the activity of Vec set at 1. n = 7 independent experiments. * p < 0.05 as compared to Vec by Student’s **E.** HEK293T/17 cells were transiently transfected with Fam64a expression vector (Fam), GR expression vector (GR), or control empty vector (Vec), and one of the three reporter constructs (ⅰ–ⅲ). Cells were treated with dexamethasone (Dex) at 1 µM for 24 h. Luciferase activity of each reporter construct was normalized to that of control reporter construct, and was expressed as the activity of Vec set at 1. n = 4 independent experiments. ** p < 0.01, *** p < 0.001 between the indicated groups by One-way ANOVA with Tukey’s post hoc test. Error bar = SEM **F.** Primary CMs were isolated from fetal hearts and transduced with baculovirus expressing Fam64a (Fam) or control empty vector (Vec) in the absence or the presence of dexamethasone (Dex) treatment at 1 µM for 24 h. Total RNA was extracted, reverse-transcribed, and subjected to qPCR analysis for *Klf15*. Data were expressed as *Klf15* mRNA expression of the vector (Vec) group set at 1. n = 3 independent experiments. In each experiment, 5–10 fetal hearts were pooled and used for the isolation of CMs. * p < 0.05, and ** p < 0.01 between the indicated groups by Student’s two-tailed unpaired t-test. Error bar = SEM.

Because Fam64a could be a putative transcriptional repressor (Archangelo et al., 2006, 2013), we tested whether Fam64a inhibits GR-mediated transcriptional activation of Klf15 using luciferase reporter assay in HEK293T/17 cells. Three reporter constructs on the human KLF15 locus was used (Figure 6C). Construct (ⅰ) contained a common promoter sequence upstream of the first exon. Construct (ⅱ) and construct (ⅲ) contained, respectively, three of the four and the remaining GR binding sites previously reported (Asada et al., 2011; Sasse et al., 2013). At baseline, in the absence of exogenous induction of GR signaling, a tendency toward a repressed activity was observed in all three constructs by Fam64a overexpression (Figure 6D). The repression was most strongly and significantly observed in the construct (ⅱ), which contains the majority of the GR binding sites (Figure 6C, D). We observed a similar repression in construct (ⅱ) in the presence of exogenous induction of GR signaling by GR overexpression and dexamethasone treatment (Figure 6E). We corroborated these findings using primary cultures of isolated CMs to show that Fam64a repressed Klf15 mRNA expression in the absence or presence of exogenous GR induction by dexamethasone (Figure 6F). Dexamethasone-induced activation of Klf15 was completely blocked by Fam64a. These data indicate that Fam64a inhibits Klf15 activity at least in part by GR-mediated transcriptional regulation through action on the previously described GR binding sites.

### Transient induction of Fam64a in cryoinjured WT hearts improves functional recovery with augmented CM cell cycle activation

On the basis of the findings that long-term induction of Fam64a in TG mice hearts exacerbates cardiac function, we next tested whether transient induction of Fam64a upon injury is beneficial. We used the protocol with direct intramyocardial injection of mRNA encoding Fam64a-FLAG or control EGFP immediately after cryoinjury in WT adult hearts (modified from Kaur and Zangi, 2020; Strungs et al., 2013). This protocol ensured the expression of the Fam64a-FLAG fusion protein in the CM nuclei (Figure 7A). The estimation by qPCR analysis revealed a 765 ± 251-fold (mean ± SEM, n = 3 mice) increase in the expression of the delivered mRNA in comparison to the non-injected controls. While cardiac contractile function was seriously damaged in both groups immediately after cryoinjury, the mice receiving Fam64a-FLAG mRNA showed progressively improved functional recovery with minimum left ventricular dilation over a follow-up period of 5 weeks in comparison to those receiving control EGFP mRNA (Figure 7B–E). The assessment of heart sections containing a cryoinjured area at 5 weeks post-injury demonstrated greater proliferative capacity in the Fam64a-FLAG group than the EGFP group, as indicated by increased positivity for Ki67 (Figure 7F). These data show that transient induction of Fam64a upon injury improved heart function with augmented CM cell cycle activation.

**Figure 7.**
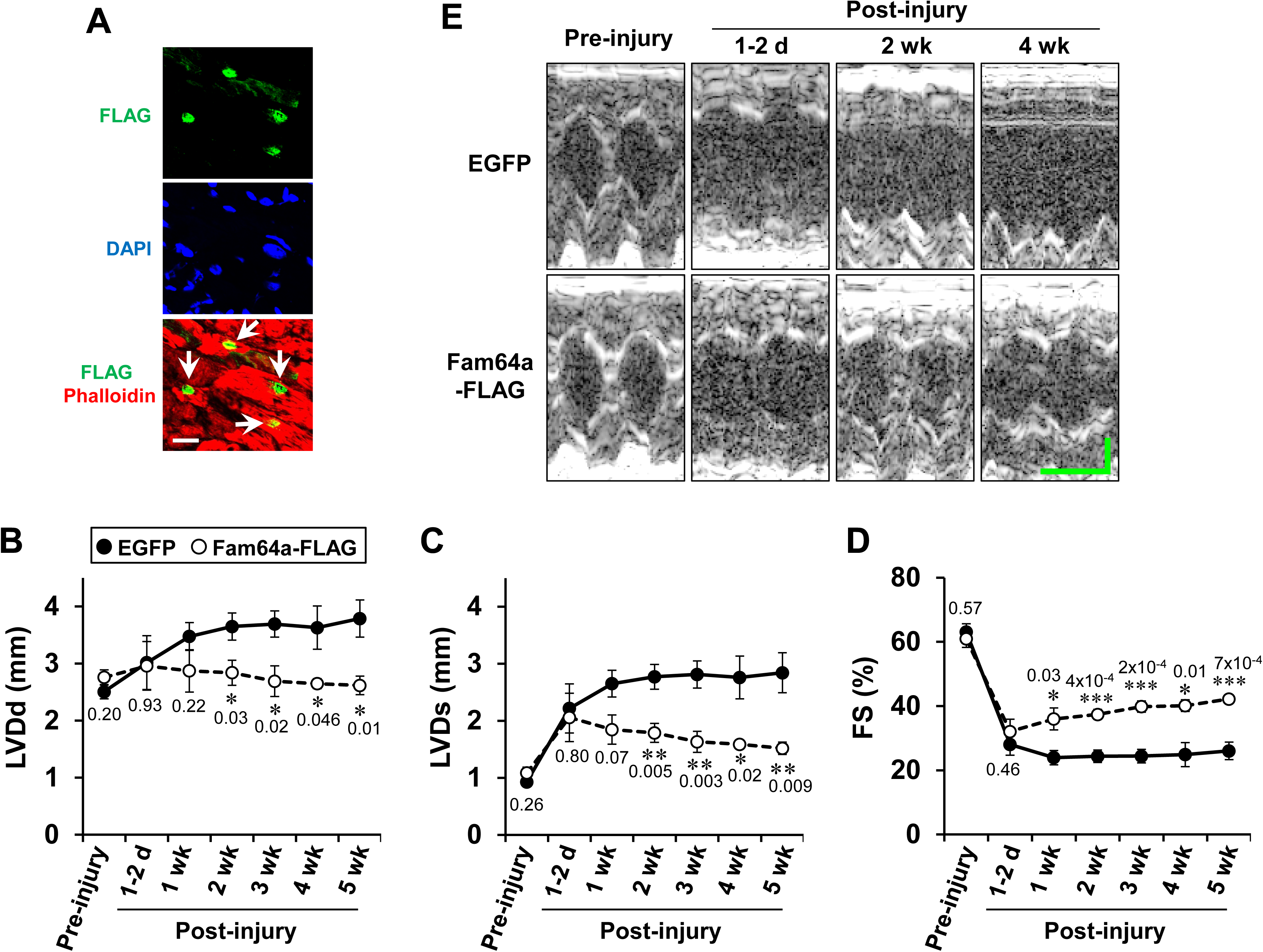
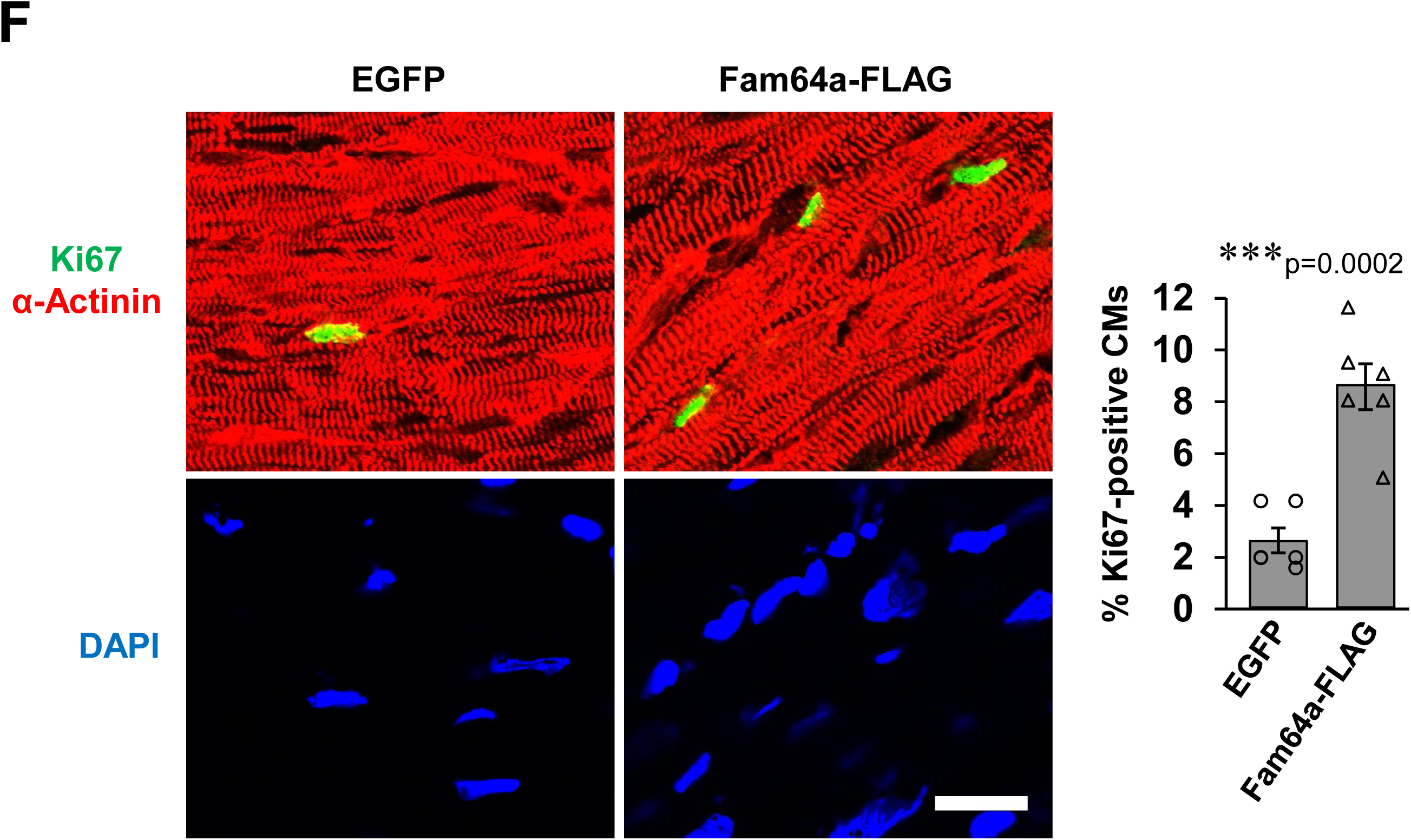
Transient induction of Fam64a in cryoinjured WT hearts improves functional recovery with augmented CM cell cycle activation. **A.** The expression of Fam64a-FLAG fusion protein was detected by immunofluorescence using anti-FLAG antibody in heart sections from mice receiving Fam64a-FLAG mRNA. The expressed protein was confirmed to localize in the CM nuclei, in the same location as an endogenous protein. Counterstaining for phalloidin and DAPI was performed. Scale bar = 20 µm **B–E.** Cardiac function was evaluated in mice receiving mRNA for EGFP (filled circle) or Fam64a-FLAG (open circle) over the course of the experiments. Left ventricular internal diameter at end diastole (LVDd, **B**) and end systole (LVDs, **C**) were measured by two-dimensional transthoracic M-mode echocardiography. Fractional shortening (FS, **D**) was calculated as ([LVDd– LVDs]/LVDd)×100 (%), and was used as an index of cardiac contractile function. Representative tracings were shown in **E** (Horizontal scale bar = 100 ms, vertical scale bar = 1 mm). Whereas cardiac function was severely declined in both groups immediately after cryoinjury, mice receiving Fam64a-FLAG mRNA showed progressively improved functional recovery with minimum left ventricular dilation over a follow-up period of 5 weeks when compared to those receiving control EGFP mRNA. n = 4–6 mice in EGFP group and 3–6 mice in Fam64a-FLAG group. * p < 0.05, ** p < 0.01, *** p < 0.001 as compared to EGFP group at the same stage by Student’s two-tailed unpaired t-test. Exact p values were shown on the graph. Error bar = SEM. **F.** Immunofluorescence for Ki67 (green) observed in sarcomeric α-actinin (red, as a CM marker) and DAPI (blue) in heart sections containing a cryoinjured area at 5 weeks post-injury from mice that had received mRNA for EGFP or Fam64a-FLAG. Quantitative analysis for the percentage of Ki67-positive CMs was shown on the right. n = 6 mice per group. For each mouse, 50–150 CMs were counted. *** p < 0.001 as compared to EGFP group by Student’s two-tailed unpaired t-test. Error bar = SEM. Scale bar = 20 µm.

## Discussion

In principle, CM cell division requires preceding dedifferentiation that accompanies the breakdown of sarcomere structures and calcium dysregulation (D’Uva et al., 2015; Zhu et al 2020), thus resulting in a depressed contractile property. Little attention has been paid to this intrinsic dilemma, which has caused cardiac dysfunction in several settings that aimed at promoting CM proliferation in the long-term (Gabisonia et al., 2019; Ikeda et al., 2019; Kubin et al., 2011).

To understand the mechanisms of this dilemma and to find the way to overcome it, we explored the feasibility of optimizing the induction protocol of the cell cycle promoter by utilizing Fam64a as a model. First, in the TG mice maintaining long-term CM-specific postnatal expression of Fam64a, we saw cardiac dysfunction characterized by the FGP-mediated sustained CM dedifferentiation and the rhythm disturbance, despite an enhancement of CM proliferation, which was reminiscent of the dilemma.

Mechanistic analysis revealed that Fam64a inhibited Klf15 activity (Figure 6). Klf15 drives CM differentiation by repressing the FGP and establishing CM rhythmic activity (Fisch et al., 2007; Jeyaraj et al., 2012a; Leenders et al., 2010, 2012; Zhang et al., 2015). Rhythmic activity of an organism is tightly coupled to cellular differentiation. The circadian clock is absent in undifferentiated cells, such as zygotes and early embryos, and is gradually established during differentiation (Umemura et al., 2017; Yagita et al., 2010). The established rhythmicity is abolished when differentiated cells are reprogrammed to regain pluripotency (Yagita et al., 2010). Therefore, overexpressing Fam64a would direct postnatal CMs toward undifferentiated states by repressing Klf15, thereby leading to dysregulation of the cardiac circadian machinery (Figure 5 and Figure 5–figure supplement 1). A previous study found that Klf15-deficient mice show perturbed CM rhythmic activity and are susceptible to ventricular arrhythmias, similarly to the effects seen in Fam64a TG mice (Figure 5C–F), which are considered to reflect suppressed KChIP2 activity (Jeyaraj et al, 2012a). KChIP2 is a critical subunit for generating the fast transient outward K^+^ current (I_to,f_) in the early repolarization phase (Kuo et al., 2001), and it augments subsequent Ca^2+^ influx in CMs (Cordeiro et al., 2012; Thomsen et al., 2009). We found severely depressed transcript levels of KChIP2 (Figure 6A) and impaired Ca^2+^ transients (Figure 3A–C) in TG mice. Suppression of K^+^ channel genes (Figure 4–figure supplement 1) would also account for the aberrant CM electrical activity (Figure 5C–F). These data suggest that long-term induction of Fam64a evoked an FGP-driven dedifferentiation through the Klf15-KChIP2 axis, thereby disrupting CM rhythmic activity and contributing to cardiac dysfunction. Thus, we propose that Fam64a is not merely a cell cycle promoter; rather, it has a previously unknown function in directing CMs toward immature undifferentiated states through inhibition of Klf15. These data also show that abnormal CM rhythmic activity is a useful indicator of dedifferentiation-mediated cardiac dysfunction, as well as sarcomere disassembly and calcium dysregulation.

We demonstrated that GR could, at least in part, mediate the inhibitory effect of Fam64a on Klf15 at the transcriptional level (Figure 6B–F). We identified Trim28 (tripartite motif-containing 28) as another interacting partner of Fam64a in CMs (Figure 6–figure supplement 1). Trim28 is a known coactivator of GR (Chang et al., 1998). Moreover, the NuRD (nucleosome remodeling and deacetylase) complex, which is implicated in gene repression (Denslow and Wade, 2007), interacts with both Fam64a (Zhao et al., 2008) and Trim28 (Schultz et al., 2001). Therefore, an important remaining challenge is to clarify the mechanism of how protein complexes comprising Fam64a, GR, and Trim28, coupled with the NuRD complex, cooperatively repress Klf15 transcription.

Mechanisms other than the Fam64a-Klf15 axis also mediate CM dedifferentiation. For example, oncostatin M has been identified as a major mediator of CM dedifferentiation (Kubin et al., 2011). Hippo deficiency and resulting activation of the Yap (Yes-associated protein)-transcriptional enhancer factor (Tead1) pathway form a positive feedback loop with oncostatin M to induce CM dedifferentiation (Ikeda et al., 2019). The potential for molecular crosstalk between these pathways and Fam64a-Klf15 axis awaits further investigations.

To overcome the intrinsic dilemma, CM redifferentiation following transient dedifferentiation and cell division is obligatory (Wang et al., 2017). In Fam64a TG mice, long-term induction caused sustained dedifferentiation, potentially inhibiting redifferentiation. This led to cardiac dysfunction even though proliferation was enhanced. In contrast, transient induction of Fam64a in the cryoinjured hearts of WT adult mice showed progressively improved functional recovery with minimum left ventricular dilation over a follow-up period of 5 weeks with augmented CM cell cycle activation (Figure 7). These data emphasize the need to optimize the intensity and duration of the stimulant to avoid excessive CM dedifferentiation. Regarding Fam64a, simultaneous activation of APC/C (anaphase-promoting complex/cyclosome), which targets Fam64a for degradation during each cell cycle (Zhao et al., 2008), deserves investigation. A recent study described the possibility for activating CM redifferentiation by showing that Ca^2+^ propagation into dedifferentiated CMs from neighboring CMs via connexin 43 gap junction provokes redifferentiation by activating calcineurin/NFAT signaling (Wang et al., 2017). In Fam64a TG mice, decreased levels of connexin 43 (Figure 4C) could thus account for the impairment of redifferentiation. Rescuing the damaged gap junction using an ischemia-resistant connexin 43 mutant (Wang et al., 2017) could facilitate CM redifferentiation.

In summary, we propose a previously unknown function of Fam64a in directing CMs toward undifferentiated immature states through inhibition of Klf15, in addition to the role as a cell cycle promoter. More importantly, our data indicate that optimizing the induction protocol, e.g., the intensity and duration, of the cell cycle stimulant, particularly to avoid excessive CM dedifferentiation is a promising option to overcome the intrinsic dilemma. Further fine-tuning of the protocol in response to various types of cardiac injury with varying degree of severity in mice and other large mammals, holds great promise for achieving successful heart regeneration, ultimately contributing to develop future regenerative therapies of the human heart.

## Acknowledgments

We are grateful to Takashi Murayama (Juntendo University, Japan) for establishing baculovirus-mediated protein expression system, and Nobuhisa Iwachido (Kawasaki Medical School, Japan) for expert technical assistance in H&E staining. This work was supported by JSPS KAKENHI Grant Number 17H02092, 18K19943, 17H06272, and 20H04521 to K.H., A.H., Y.Ujihara., and S.M. from the Ministry of Education, Culture, Sports, Science, and Technology of Japan, and was also supported by grants from Kawasaki Medical School.

## Author contributions

KH designed and performed the majority of the experiments, and wrote the manuscript; AK, MO, and MK performed biochemical and molecular biology experiments, and analyzed data; RN performed mass spectrometry experiments and analyzed data; YUsui performed cryoinjury experiments and analyzed data; YUjihara performed experiments using isolated CMs and analyzed data; AH contributed to molecular cloning, plasmid construction, and baculovirus production; SM supervised the study and contributed to manuscript preparation. All authors critically read and approved the manuscript.

## Declaration of interests

The authors declare no competing interests.

## Methods

This study was performed in strict accordance with the recommendations of the Institutional Animal Care and Use Committee at the Kawasaki Medical School. All of the animals were handled according to approved institutional protocols of the Kawasaki Medical School, and every effort was made to minimize suffering. All experiments were performed in accordance with the relevant guidelines and regulations of the Kawasaki Medical School.

### Mice

Mice with a C57BL/6N background were housed in a temperature-controlled room under a 12-h light:12-h dark cycle conditions and were fed a standard chow diet and water ad libitum. CM-specific Fam64a TG mice were generated as follows: the murine Fam64a sequence with a C terminal FLAG tag was cloned downstream of the alpha myosin heavy chain promoter (Figure 1–figure supplement 1). The alpha myosin heavy chain/puro rex/neo was a gift from Mark Mercola (Addgene plasmid #21230; http://n2t.net/addgene:21230) (Kita-Matsuo et al., 2009). This transgene construct was purified, linearized, and injected into fertilized oocytes from C57BL/6N background mice (Transgenic Inc, Japan). The resulting pups were genotyped by PCR using genomic tail DNA and seven founder lines were established. Among these, two lines expressing a sufficient amount of the transgene both at mRNA and protein levels were selected and used for subsequent experiments (Figure 1–figure supplement 1). Wildtype (WT) mice with the same background were used for comparison.

### Immunofluorescence

Frozen heart sections embedded in OCT compound (Tissue-Tek®; Sakura, UAE) were cut into 8 µm sections with a cryostat (Leica, Germany), permeabilized, blocked with Blocking-One (Nacalai Tesque, Japan), and labelled with primary antibodies, followed by fluorochrome-conjugated secondary antibodies. The sections were also counterstained with DAPI (nuclei) and phalloidin (F-actin). Sections were covered with a fluorescence mounting medium (Dako, USA) and examined using a confocal scanning system mounted on a IX81 inverted microscope (FV-1000, Olympus, Japan), as previously described (Hashimoto et al., 2018). The primary antibodies used were for Ki67 (clone SP6; Abcam, UK), phospho-histone H3 at Ser-10 (EMD Millipore, USA), sarcomeric α-actinin (A7811, Sigma-Aldrich, USA), and the FLAG tag (F1804, Sigma-Aldrich).

### Western blotting

Heart tissues were collected from the mice and snap frozen in liquid nitrogen, minced, and homogenized using a Kinematica™ Polytron™ homogenizer (PT1600E; Fisher Scientific, USA) in M-PER extraction buffer (Thermo-Fisher, USA) in the presence of a protease inhibitor cocktail (Thermo-Fisher). Lysates were centrifuged at 14,000×g and supernatants were used as the whole protein extract. After quantifying the protein yield, equal amounts of protein were separated by SDS-PAGE (Mini-PROTEAN® TGX; Bio-Rad, USA), transferred onto PVDF membranes (GE Healthcare, USA), blocked with 5% nonfat milk, probed with primary antibodies followed by secondary horseradish peroxidase (HRP)-conjugated IgG (GE Healthcare), and finally visualized by enhanced chemiluminescence (Western Lightning ECL-Pro; PerkinElmer, USA) using a LAS4000mini luminescent image analyzer (GE Healthcare), as previously described (Hashimoto et al., 2017). Primary antibodies used were for Trim28 (#4124, Cell Signaling Technology, USA), GR (sc-393232, Santa Cruz Biotechnology, USA), the FLAG tag (F1804, Sigma-Aldrich), and Fam64a. The Fam64a antibody was raised against a synthetic peptide corresponding to residues 154–172 of mouse Fam64a (CRLSGQMGPHAHRRQRLRRE).

### Immunoprecipitation and mass spectrometry

We identified the interacting partners of Fam64a using immunoprecipitation against the FLAG peptide that was expressed as a C-terminal tag of the overexpressed Fam64a protein in TG mice hearts, followed by mass spectrometry analysis (n = 2 biological replicates). Immunoprecipitates from WT mice hearts were used as a negative control. Heart tissues were freshly isolated from WT and TG mice, minced, and homogenized using a Kinematica™ Polytron™ homogenizer (Fisher Scientific) in IP lysis buffer (Thermo-Fisher) or cytoplasmic extraction reagent I & II (Thermo-Fisher) in the presence of a protease inhibitor cocktail (Thermo-Fisher). After centrifugation and protein quantification, the lysates were subjected to immunoprecipitation using the EZview Red Anti-FLAG M2 affinity gel system (F2426, Sigma-Aldrich) according to the manufacturer’s instructions. Elution of the immunoprecipitates was performed with 3× FLAG peptide (F4799, Sigma-Aldrich). The immunoprecipitation procedure was validated by western blotting using FLAG and Fam64a antibodies, which correctly detected the Fam64a-FLAG fusion protein in TG, but not in WT, mouse heart lysates (Figure 6–figure supplement 1).

#### LC-MS/MS ANALYSIS

In-solution digestion and nano flow-liquid chromatography tandem mass spectrometry were performed as described previously (Oya et al., 2019), with some modifications. In brief, the eluted proteins were digested with 10 μg/mL modified trypsin (Sequencing grade, Promega, USA) at 37 °C for 16 h. The digested peptides were desalted with in-house made C18 Stage-tips, dried under a vacuum, and dissolved in 2% acetonitrile containing 0.1% trifluoroacetic acid. The peptide mixtures were then fractionated by C18 reverse-phase chromatography (3 μm, ID 0.075 mm x 150 mm, CERI). The peptides were eluted at a flow rate of 300 nL/min with a linear gradient of 5–35% solvent B over 90 min.

#### DATABASE SEARCHING

The raw files were searched against the *Mus musculus* dataset (Uniprot Proteome ID UP000000589 2019.06.11 downloaded, 55,197 sequences; 22,986,518 residues) combined with the FLAG-tagged Fam64a sequence and the common Repository of Adventitious Proteins (cRAP, ftp://ftp.thegpm.org/fasta/cRAP) using MASCOT version 2.6 (Matrix Science) via Proteome discoverer 2.2 (Thermo-Fisher), with a false discovery rate (FDR) set at 0.01. Carbamidomethylation of cysteine was set as a fixed modification. Oxidation of methionine and acetylation of protein N-termini were set as variable modifications. The number of missed cleavage sites was set as 2.

#### CRITERIA FOR PROTEIN IDENTIFICATION

Scaffold (version Scaffold_4.10.0, Proteome Software Inc., Portland, OR) was used to validate the MS/MS-based peptide and protein identifications. Peptide identifications were accepted if they exceeded specific database search engine thresholds. Protein identifications were accepted if they contained at least 2 identified peptides. Proteins that contained similar peptides and could not be differentiated based on MS/MS analysis alone were grouped to satisfy the principles of parsimony. Proteins sharing significant peptide evidence were grouped into clusters.

In two biologically independent experiments, 335 and 796 proteins were detected under the threshold setting in Scaffold software, as follows: protein threshold of 1.0% FDR, peptide threshold of 0.1% FDR, and Min # peptides = 2. Proteins detected only in TG samples, but not in WT samples, were considered as candidate interacting partners of Fam64a, and the interaction of those proteins with Fam64a in heart tissues was subsequently confirmed by immunoprecipitation and western blotting using specific antibodies. All mass spectrometry data have been deposited in ProteomeXchange Consortium via jPOST, with the dataset identifiers PXD020570 and JPST000921 (Preview code for reviewers https://repository.jpostdb.org/preview/497638005f1cd5ace27c1, Access key: 6340).

### Quantitative PCR (qPCR)

Heart tissues were collected from mice, cut into small pieces, and immediately immersed in RNAlater® Stabilization Reagent (Qiagen, Germany). The stabilized tissues were homogenized with a Kinematica™ Polytron™ homogenizer (Fisher Scientific), and total RNA was isolated using the ISOGEN or ISOGEN-II systems (Nippon Gene, Japan). For cultured CMs, harvested cell pellets were processed similarly to heart tissues but without the use of the homogenizer. After assessing RNA yield and quality using a NanoDrop One spectrophotometer (Thermo-Fisher), the RNA samples were reverse-transcribed with PrimeScrip RT Master Mix (TaKaRa Bio, Japan), and quantitative real-time PCR was performed using TaqMan® Fast Advanced Master Mix in a StepOnePlus™ real-time PCR system (Applied Biosystems, USA). Quantification of each mRNA was carried out with *Actb* or *Ubc* as reference genes, using the ΔΔC_T_ method, as previously described (Hashimoto et al., 2018).

### Luciferase reporter assay

Three reporter constructs spanning the promoter region of human KLF15 locus were used (Figure 6C): construct (ⅰ): −694/+228, construct (ⅱ): +1066/+1965, and construct (ⅲ): +9444/+10643, where the number indicates the genomic position relative to the transcription start site. The sequence of the construct (ⅰ) was derived from the LightSwitch™ Promoter Reporter GoClone™ (SwitchGear Genomics, USA). The expression vectors were a Fam64a expression vector, GR expression vector, or control empty vector (pFastBac1-VSVG-CMV-WPRE; Hashimoto et al., 2017). HEK293T/17 cells (ATCC® CRL-11268™) were maintained in DMEM with 5% FBS under standard conditions at 37 °C with 5% CO_2_. Cells were plated onto 96-well plates coated with fibronectin, and transient transfection of the reporter construct and the expression vector was carried out using Lipofectamine® 2000 (Thermo-Fisher) on the following day. The amount of plasmid used per well was 3 ng for each expression vector / 50 ng for each reporter construct. The control empty vector was used to equalize the total amount of DNA for each transfection. The expression of the Fam64a protein was confirmed by western blotting. Cells were treated with dexamethasone (Dex) at 1 µM for 24 h. Luciferase activity was measured on the next day using the LightSwitch™ luciferase assay system (SwitchGear Genomics) as per the manufacturer’s protocol, as previously described (Hashimoto et al., 2017). The luciferase activity of each reporter construct was normalized to that of the control reporter construct (pLightSwitch_Prom) and was expressed as the activity of the control empty vector set at 1.

### Histology

Heart tissues were collected from mice, fixed in 4% paraformaldehyde, embedded in paraffin, and vertically sectioned at a thickness of 3 μm. Hematoxylin-eosin (H&E) staining was performed according to standard procedures. Stained sections were observed with a light microscope (BZ-X710, Keyence, Japan).

### Echocardiography

Two-dimensional transthoracic echocardiography was performed to evaluate cardiac function using an Aplio 300 system with a 14-MHz transducer (Toshiba Medical System, Japan), as previously described (Ujihara et al., 2016). Mice were initially anesthetized with 2% sevoflurane and then maintained on 1% sevoflurane during examinations. M-mode tracings were used to measure the left ventricular internal diameter at end diastole (LVDd) and end systole (LVDs). Fractional shortening (FS) was calculated as ([LVDd–LVDs]/LVDd)×100 (%), and was used as an index of cardiac contractile function.

### Locomotor activity measurement

The locomotor activity of mice was monitored using an infrared motion detector (Actimo-100, Shinfactory, Japan), which consists of a free moving space (30 × 20 cm^2^) with a side wall equipped with photosensors at 2 cm intervals to scan animal movement as described (Kurokawa et al., 2011). Activity counts accumulated over a 1 h period were measured for a total of 4 days in a 12 h light:12 h dark cycle (lights on at 8 am). Total activity counts during the daytime (8 am to 8 pm) and nighttime (8 pm to 8 am) were considered to reflect the locomotor activity in each phase. During the nighttime, we found that the most mice showed characteristic biphasic patterns of locomotor activity, i.e., the first peak during the time period from 8 pm to 2 am, and the second peak during the time period from 2 am to 8 am (typical example shown in Figure 5–figure supplement 1). Thus, the peak activity counts in each phase were used as a measure of the locomotor activity during nighttime.

### ECG telemetry

Mice were anesthetized with 3% sevoflurane and implanted with a telemetry transmitter device subcutaneously (PhysiolTel® HD-X11, Data Sciences International, USA). Two ECG leads were secured at the apex of the heart and the right acromion. The ECG traces of the conscious mice were continuously recorded with a scheduled sampling (10 s every 1 min) using Dataquest ART 4.0 software (Data Sciences International) for a total of 8 days in a 12 h light:12 h dark cycle (lights on at 8 am). The heart rate was calculated by digital tracking of the ECG RR intervals using the Dataquest software, and averaged over 12 h during daytime (8 am to 8 pm) and nighttime (8 pm to 8 am). Frequency of premature ventricular contraction was examined over the two representative periods (10 min each, 5000–6000 beats) per animal, which were selected in the middle of the 8-day measurement period (Day 4 and Day 5).

### Synthesis of modified mRNA

Murine Fam64a-FLAG or EGFP sequence downstream of T7 promoter was cloned into pGEMHE, and was PCR amplified using a primer set (3’-GTAAAACGACGGCCAGT-5’ and 3’-CAGGAAACAGCTATGAC-5’). This PCR product was purified using FastGene Gel/PCR Extraction Kit (Nippon Genetics, Japan), and was used as the template for modified mRNA synthesis. In vitro transcription was performed using HiScribe T7 High Yield RNA Synthesis Kit (New England Biolabs, USA) with a customized ribonucleoside blend of GTP (1.5 mM), ATP (7.5 mM), CTP (7.5mM), N1-Methylpseudo-UTP (7.5 mM, N-1081, TriLink Biotechnologies), and CleanCap® Reagent AG (6 mM, N-7113, TriLink Biotechnologies, USA) (Zangi et al., 2017; Kaur and Zangi, 2020). Following the purification of transcribed mRNA using Fast Gene RNA Premium Kit (Nippon Genetics), Poly (A) tailing reaction was performed using E. coli Poly (A) polymerase (New England Biolabs), and mRNA was re-purified with the same kit. The size and the integrity of synthesized modified mRNA was checked by agarose gel electrophoresis, and quantity was determined using a NanoDrop One spectrophotometer (Thermo Scientific).

### Transient induction of Fam64a in cryoinjured hearts using modified mRNA

Cryoinjury experiments were perfomed using protocols adapted from Strungs et al., 2013. WT male mice (13–25 weeks old) were anesthetized with a mixture of 0.3 mg/kg medetomidine, 4.0 mg/kg midazolam, and 5.0 mg/kg butorphanol via subcutaneous injection (Tashiro et al., 2020). Mice were then intubated and mechanically ventilated at 80 breaths/min with a tidal volume of 1000 μL using a rodent ventilator (SN-480-7, Shinano manufacturing, Japan). Hearts were exposed by a left thoracotomy at the fourth intercostal space, the pericardial sac was gently opened by blunt dissection, and a 3-mm-diameter metal cryoprobe, prechilled with liquid nitrogen for 40 s, was applied to the epicardial surface of left ventricular free wall near the apex for 40 s. Cryoinjured area was visually confirmed as a uniform white spot. Immediately after cryoinjury, 20 μg of modified mRNA for Fam64a-FLAG was delivered via direct intramyocardial injection near the cryoinjured spot in a total volume of 100 μL using Lipofectamine™ RNAiMAX Transfection Reagent (Thermo-Fisher) (modified from Zangi et al., 2017). Modified mRNA for EGFP was used as a negative control. The chest and skin were closed and mice were allowed to recover on a heating pad until normal respiration was obtained. In this procedure, the delivery of modified mRNA at the time of cryoinjury allowed to avoid a second surgery later, which contributed to reduce mortality rate. Echocardiography was performed before and after cryoinjury to assess cardiac function over the course of experiments as described in the aforementioned section. At the end of the experiments (5 weeks after cryoinjury), mice were sacrificed, and frozen tissue sections containing a cryoinjured region were assessed for the expression of cell cycle marker Ki67 by immunofluorescent analysis, as described in the aforementioned section.

### RNA-seq

Heart tissues were collected from mice, cut into small pieces, and immediately immersed in RNAlater® Stabilization Reagent (Qiagen). The stabilized tissues were homogenized with a Kinematica™ Polytron™ homogenizer (Fisher Scientific), and total RNA was isolated using ISOGEN or ISOGEN-II system (Nippon Gene). After assessing RNA yield and quality using a 2100 Bioanalyzer (Agilent Technologies, USA), RNA-seq libraries were generated using the TruSeq Stranded mRNA Library Prep Kit (Illumina, USA). The quality of the libraries was checked using the 2200 TapeStation (Agilent Technologies). Paired-end sequencing of the libraries was performed on an Illumina Hiseq 2500 platform (Hokkaido System Science, Japan). The obtained data were processed as follows: known adapters and low-quality regions of the reads were trimmed using cutadapt 1.1 (https://cutadapt.readthedocs.org/en/stable/) and Trimmomatic 0.32 (http://www.usadellab.org/cms/index.php?page=trimmomatic), respectively. The reads were mapped to the mouse reference genome (GRCm38, Release 92) using Tophat 2.0.14 (http://ccb.jhu.edu/software/tophat/index.shtml). Gene expression between samples was compared by calculating normalized expression values for each transcript as fragments per kilobase of exon model per million mapped fragments (FPKM) using Cufflinks 2.2.1 (http://cole-trapnell-lab.github.io/cufflinks/). The expression changes in TG vs. WT mice for more than 20 genes were validated by qPCR. Enrichment of genetic associations within KEGG pathways was determined by functional annotation analysis using DAVID (https://david.ncifcrf.gov/summary.jsp). In this analysis, genes upregulated > 2.0 and downregulated < 0.5 in TG relative to WT mice were used to identify the differentially regulated gene pathways. RNA-seq data have been deposited in DDBJ sequencing read archive (DRA) under the accession number DRA009818 (http://trace.ddbj.nig.ac.jp/DRASearch).

### CM isolation from aged mice

Primary CMs were isolated from the ventricles of mice at aged stages (29–32 wks), as previously described (Ujihara et al., 2019). Briefly, the heart was excised and a cannula was inserted into the aorta. Coronary perfusion was initiated with cell-isolation buffer (CIB; 130 mM NaCl, 5.4 mM KCl, 0.5 mM MgCl_2_, 0.33 mM NaH_2_PO_4_, 22 mM glucose, 50 nM/ml bovine insulin, and 25 HEPES-NaOH (pH 7.4)) containing 0.4 mM EGTA. The perfusate was changed to the enzyme solution in CIB containing 0.3 mM CaCl_2_, 1 mg/ml collagenase type II (Worthington Biochemical, USA), 0.06 mg/ml protease (Sigma-Aldrich), and 0.06 mg/ml trypsin (Sigma-Aldrich). The left ventricles were cut into small pieces and further digested in the enzyme solution for 10–15 min at 37°C by gentle agitation. In this enzyme solution, the CaCl_2_ level was increased to 0.7 mM, and 2 mg/ml BSA was supplemented. After centrifugation at 14 × g for 5 min, the pellet was resuspended in CIB containing 1.2 mM CaCl_2_ and 2 mg/ml BSA, and then incubated for 10 min at 37°C. After a further centrifugation, the cells were resuspended in Tyrode’s solution (140 mM NaCl, 5.4 mM KCl, 1.8 mM CaCl_2_, 0.5 mM MgCl_2_, 0.33 mM NaH_2_PO_4_, 11 mM glucose, 2.0 5 mM HEPES-NaOH (pH=7.4)) containing 2 mg/ml BSA.

### Cell shortening and Ca^2+^ transient measurements

Electrically evoked cell shortening and Ca^2+^ transients were determined in isolated CMs from aged mice stimulated in an electrical field using a two-platinum electrode insert connected to an isolator (SS-104J, Nihon Kohden, Japan) and a bipolar stimulator (SEN-3401, Nihon Kohden), as previously described (Katanosaka et al., 2014). The cells were monitored with a CMOS camera (ORCA flash 4.0, Hamamatsu Photonics, Japan) mounted on the side port of an inverted microscope (IX73, Olympus) with a 20× objective lens (UCplanFLN, Olympus). The Ca^2+^ transients were measured by loading isolated CMs with 5 μM Fura-2 AM (Dojindo, Japan) for 30 min, as previously described (Honda et al., 2018). The Fura-2-loaded cells were alternately excited at 340 and 380 nm using an LED illuminator (pE-340^fura^, CoolLED). The Ca^2+^ content of the sarcoplasmic reticulum (SR) was evaluated by rapidly applying 10 mM caffeine and measuring the resulting Ca^2+^ transients in isolated CMs. Data were analyzed using MetaMorph version 7.8.0.0 software (Molecular Devices, USA).

### CM isolation from fetal mice

Primary CMs were isolated from the ventricles of fetal mice at embryonic day E17, essentially as described previously (Hashimoto et al., 2018). Briefly, pregnant mice were euthanized with Sevofrane, and fetal heart ventricles were rapidly excised, cut into small pieces, and digested four times with 0.06% trypsin and 0.24 mM EDTA in PBS for 10 min at 37°C. After a 20 min culture to exclude non-CMs, the CMs were plated onto fibronectin-coated culture vessels in DMEM with 5% FBS and cultured under standard conditions at 37 °C with 5% CO_2_. In each isolation procedure, 5–10 fetal hearts were pooled and used for the isolation.

### Baculovirus-mediated protein expression

We have previously established a baculovirus-mediated protein expression system in CMs (Hashimoto et al., 2017, 2018). In this study, a baculovirus expressing the full-length mouse Fam64a or a control empty vector was used. Baculovirus was produced in Sf9 cells, as per the manufacturer’s instructions (Thermo-Fisher). For transduction to CMs, virus was added to the cells in MEM without serum. After 7 h, the cells were treated with BacMam enhancer (Invitrogen, USA) for an additional 2 h, according to the manufacturer’s protocol, to increase the transduction efficiency. The medium was then replaced with DMEM containing 5% FBS.

### Statistics

All data were expressed as mean plus or minus standard error of the mean (SEM). For comparisons between two groups, Student’s two-tailed unpaired t-test was performed using Microsoft Excel 2019 MSO (16.0.10358.20061) to determine statistical significance. For comparisons among multiple groups, one-way analysis of variance (ANOVA) was performed with Tukey’s post hoc test using SPSS statistics ver. 26 (IBM, USA). Kaplan–Meier analysis was performed using SPSS statistics ver. 26 (IBM) to estimate the survival curve of WT and TG mice, and between-group differences were analyzed with the log-rank test. p < 0.05 was considered statistically significant. Significance levels were indicated as follows: p < 0.05, ‘*’ p < 0.01, ‘**’ p < 0.001‘***’. Additional statistical information, including sample sizes and p-values for each experiment, is detailed in the figure legends.

---

**Figure 1–figure supplement 1.**
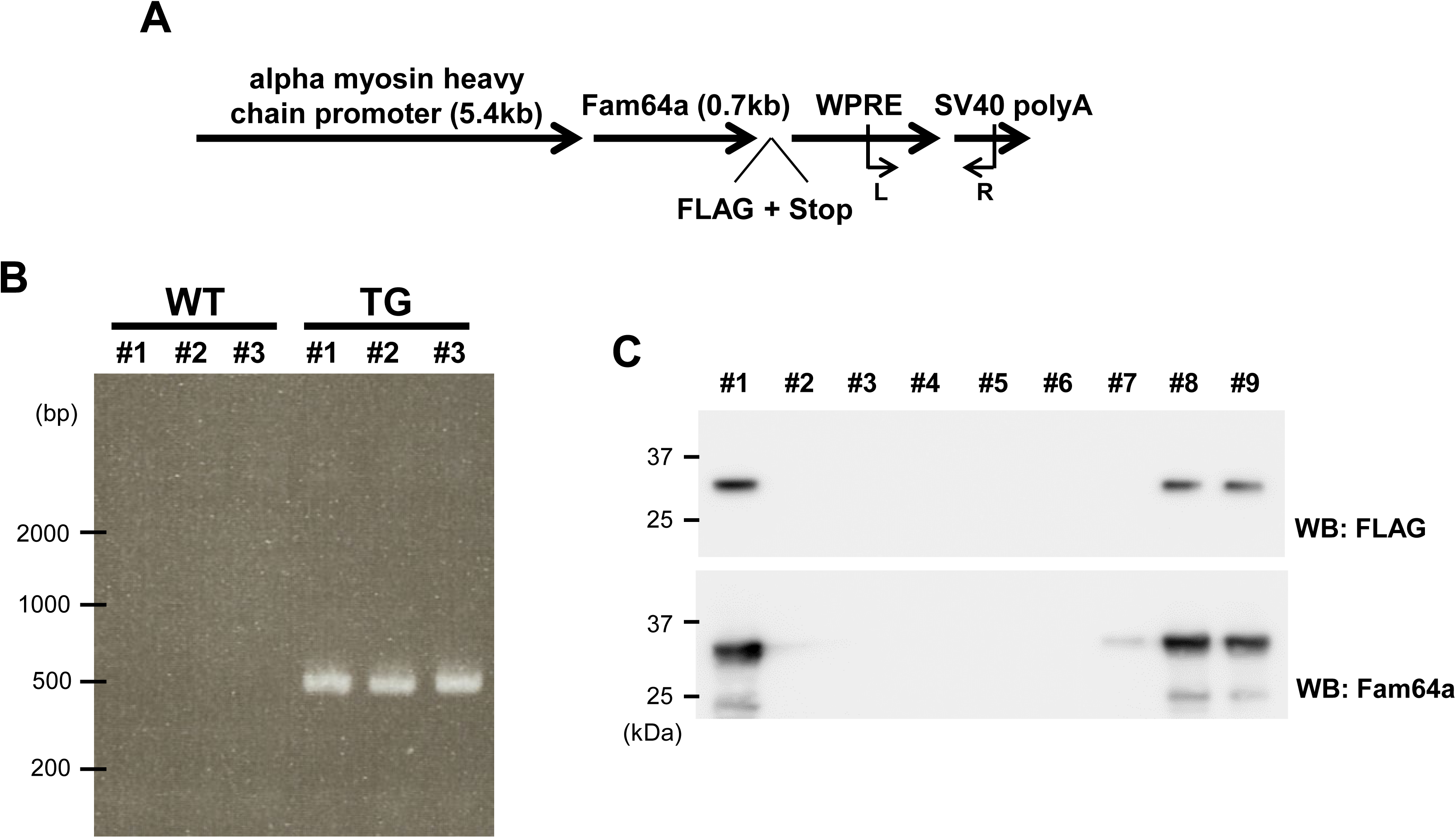
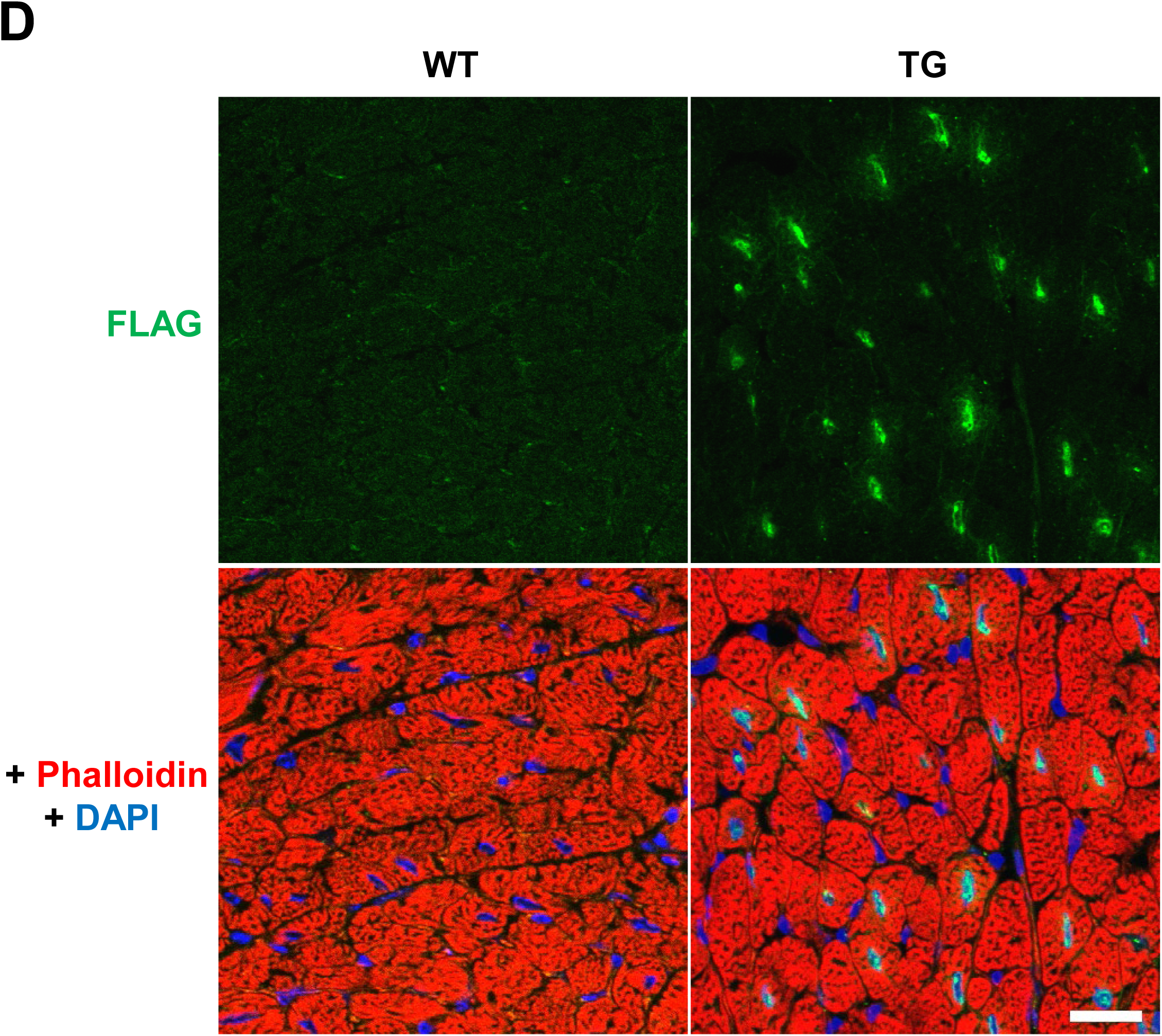
Generation of cardiomyocyte-specific Fam64a TG mice. **A.** The transgene construct containing murine Fam64a sequence with a C terminal FLAG tag cloned downstream of the alpha myosin heavy chain promoter. WPRE; Woodchuck hepatitis virus Posttranscriptional Regulatory Element for stabilizing transcribed mRNA. **B.** Representative genotyping results using genomic tail DNA from 3 WT and 3 TG mice. Positions for left (L) and right (R) primers were marked in A. **C.** Representative western blots (WB) of heart homogenates from 9 mice at F1 generation (6 wks of age) derived from 7 founder TG lines using anti-FLAG or anti-Fam64a antibody, both of which detect overexpressing Fam64a-FLAG fusion protein. Based on the amount of the protein detected, we classified 7 founder lines into 3 categories; TG-strong (2 lines, for example #1, #8, and #9), TG-medium (2 lines, for example #7), and TG-weak (3 lines, for example #2, #4, and #5). Descendants of TG-strong lines were used for subsequent experiments. **D.** Immunofluorescence for heart tissue sections from WT and TG mice at 6 wks using anti-FLAG antibody that detects overexpressing Fam64a-FLAG fusion protein. Counterstaining for phalloidin and DAPI was performed. Expressed protein was confirmed to localize in the CM nuclei, in the same location as an endogenous protein. Scale bar = 20 µm.

**Figure 3–figure supplement 1.**
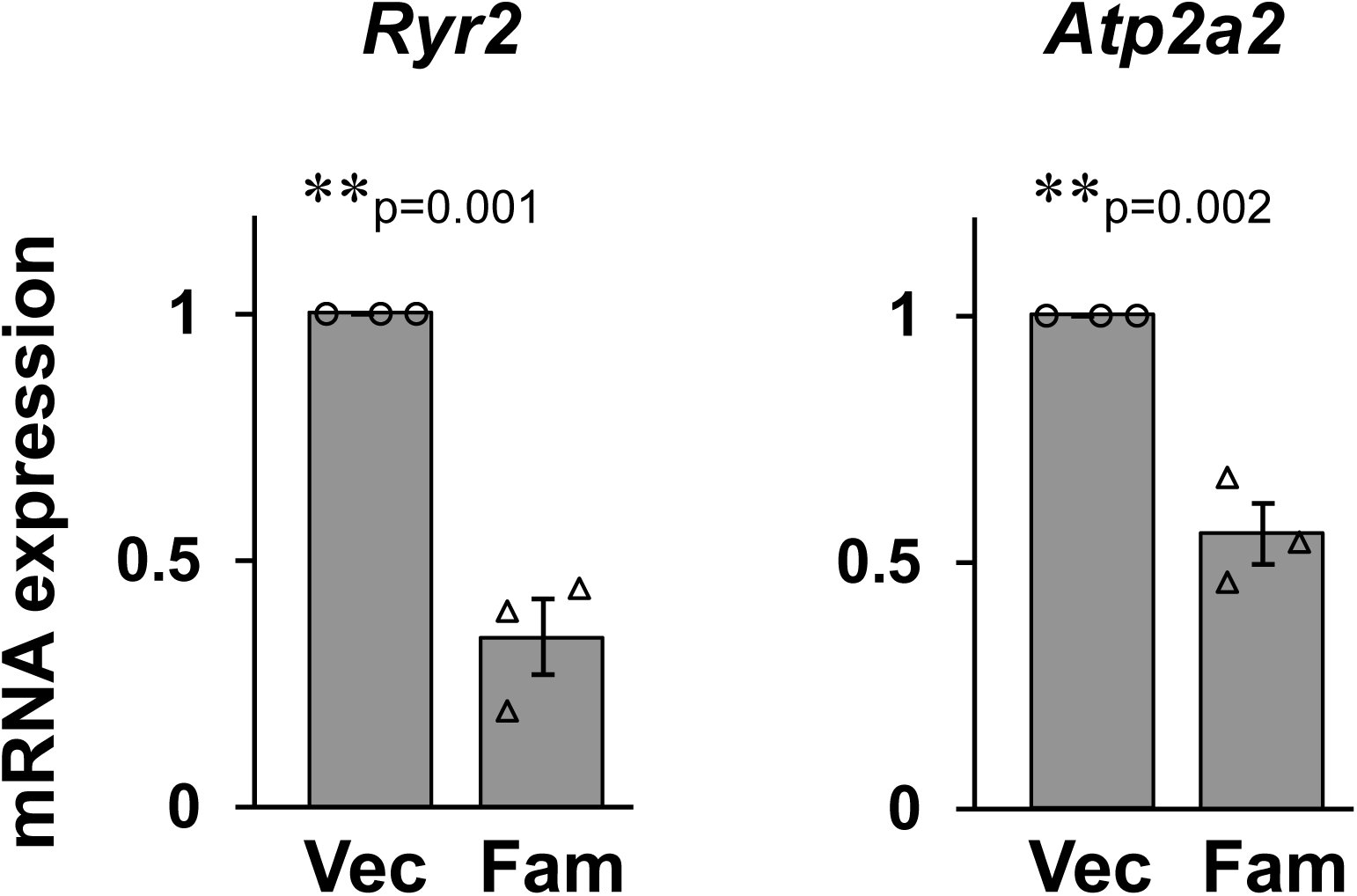
Ca^2+^ handling genes were downregulated by Fam64a overexpression in isolated CMs. Primary CMs were isolated from fetal hearts and transduced with baculovirus expressing Fam64a (Fam) or control empty vector (Vec). Total RNA was extracted, reverse-transcribed, and subjected to qPCR analysis for *Ryr2* and *Atp2a2*. Data were expressed as mRNA expression in the vector group set at 1. n = 3 independent experiments. In each experiment, 5–10 fetal hearts were pooled and used for the isolation of CMs. ** p < 0.01 as compared to Vec by Student’s two-tailed unpaired t-test.. Error bar = SEM.

**Figure 4–figure supplement 1.**
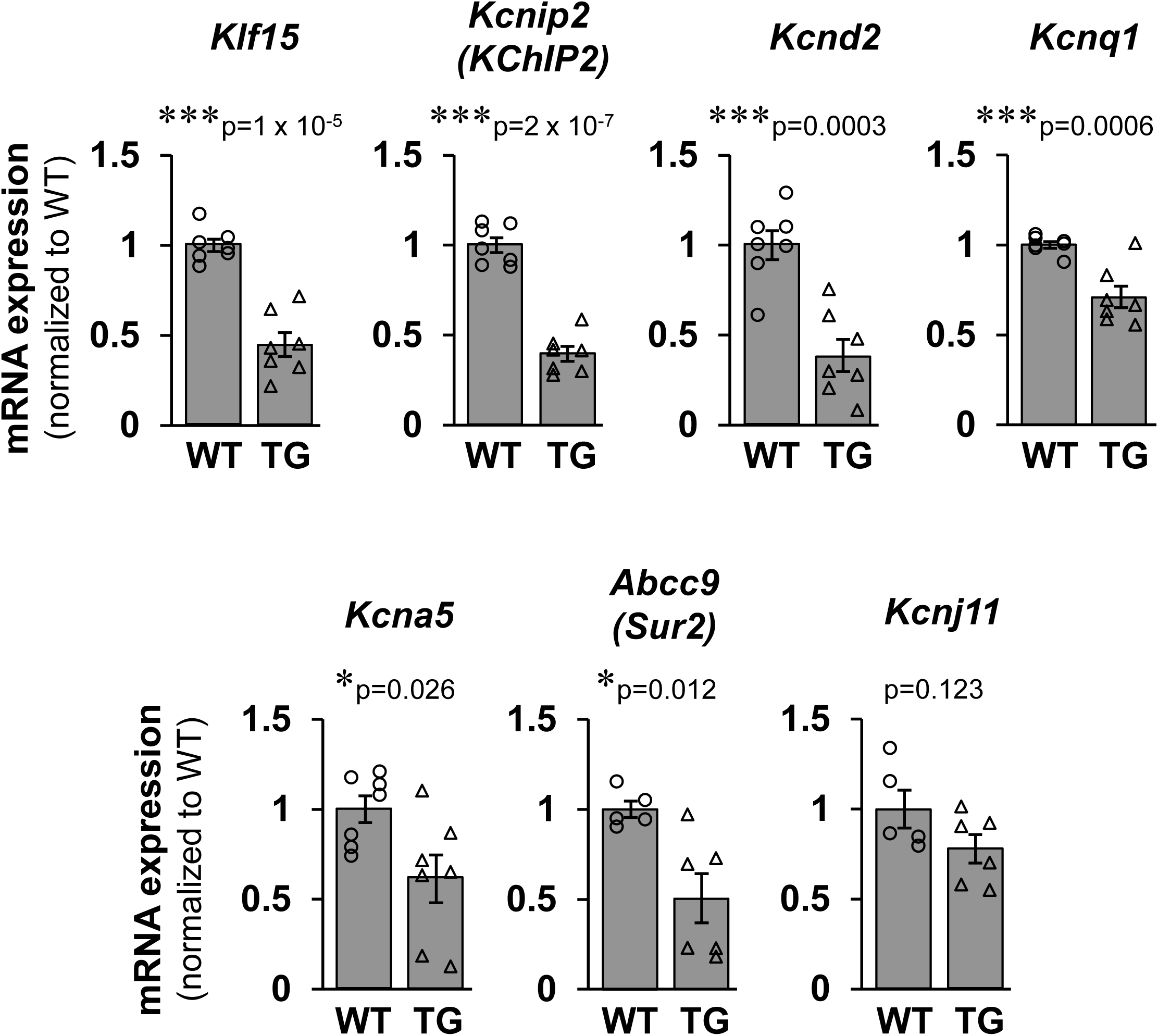
Genes for K^+^ channel subunits were consistently repressed in Fam64a TG mice. qPCR analysis of genes encoding several K^+^ channel subunits in WT and TG mice hearts. Data were shown as normalized to WT. n = 5–7 mice per group. * p < 0.05, *** p < 0.001 as compared to WT by Student’s two-tailed unpaired t-test. Error bar = SEM.

**Figure 5–figure supplement 1.**
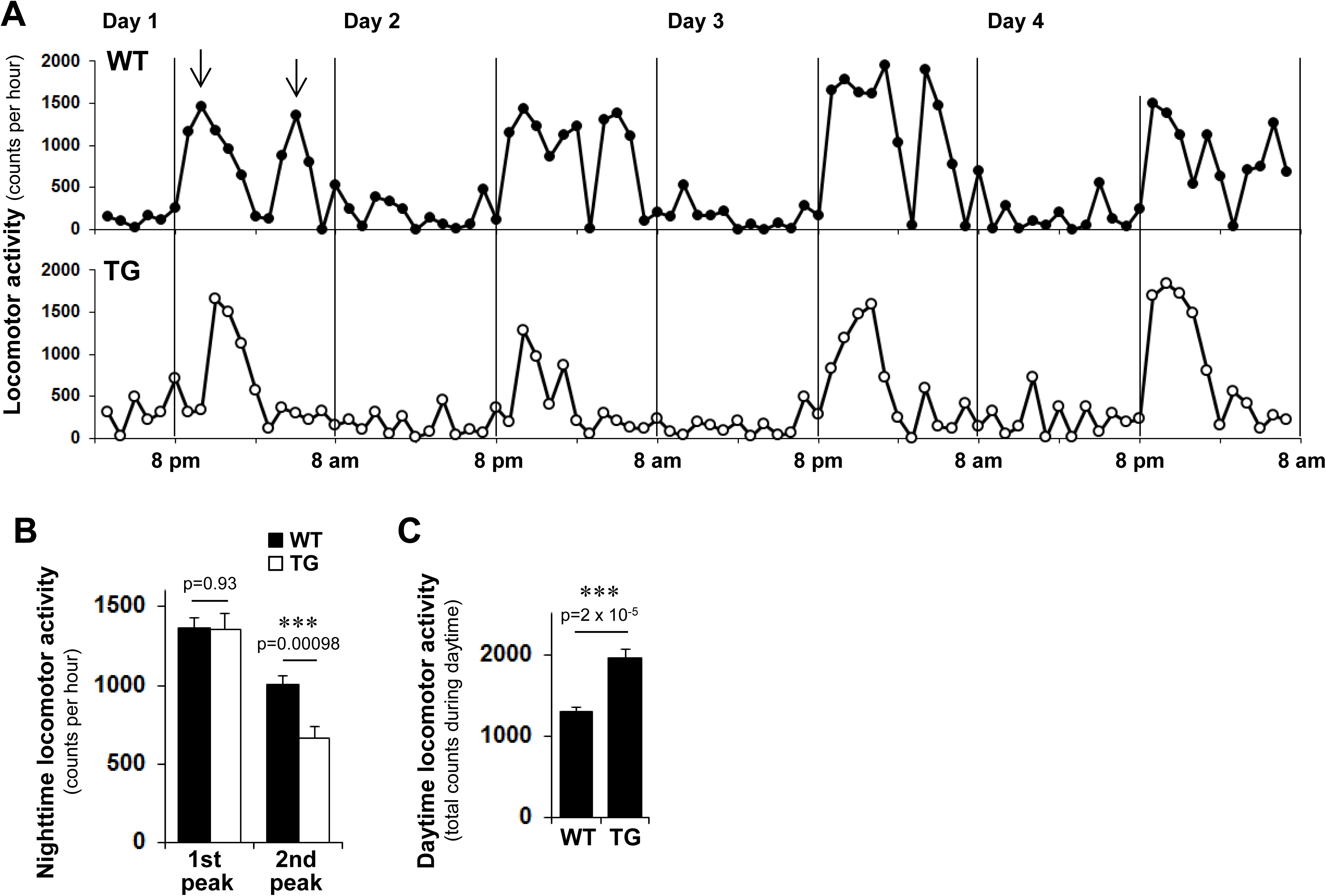
Perturbed locomotor activity in Fam64a TG mice. The locomotor activity of mice was monitored using the infrared motion detector. Activity counts accumulated over a 1 h period were measured for a total of 4 days in a 12 h light:12 h dark cycle (lights on at 8 am). **A:** Representative tracings of the locomotor activity from WT (top) and TG (bottom) mice. During the nighttime (8 pm to 8 am), the most mice showed characteristic biphasic patterns of locomotor activity, i.e., the 1^st^ peak during the time period from 8 pm to 2 am, and the 2^nd^ peak during the time period from 2 am to 8 am (arrows). **B:** The 1^st^ and the 2^nd^ peak activity during nighttime in WT (filled bar) and TG (open bar) mice. **C:** Total activity counts during daytime (8 am to 8 pm) in WT and TG mice. Data pooled for 4 days were shown. Data were analyzed from 7 WT mice and 8 TG mice at adult stages (9–16 wks of age). *** p < 0.001 as compared to WT by Student’s two-tailed unpaired t-test. Error bar = SEM.

**Figure 6–figure supplement 1.**
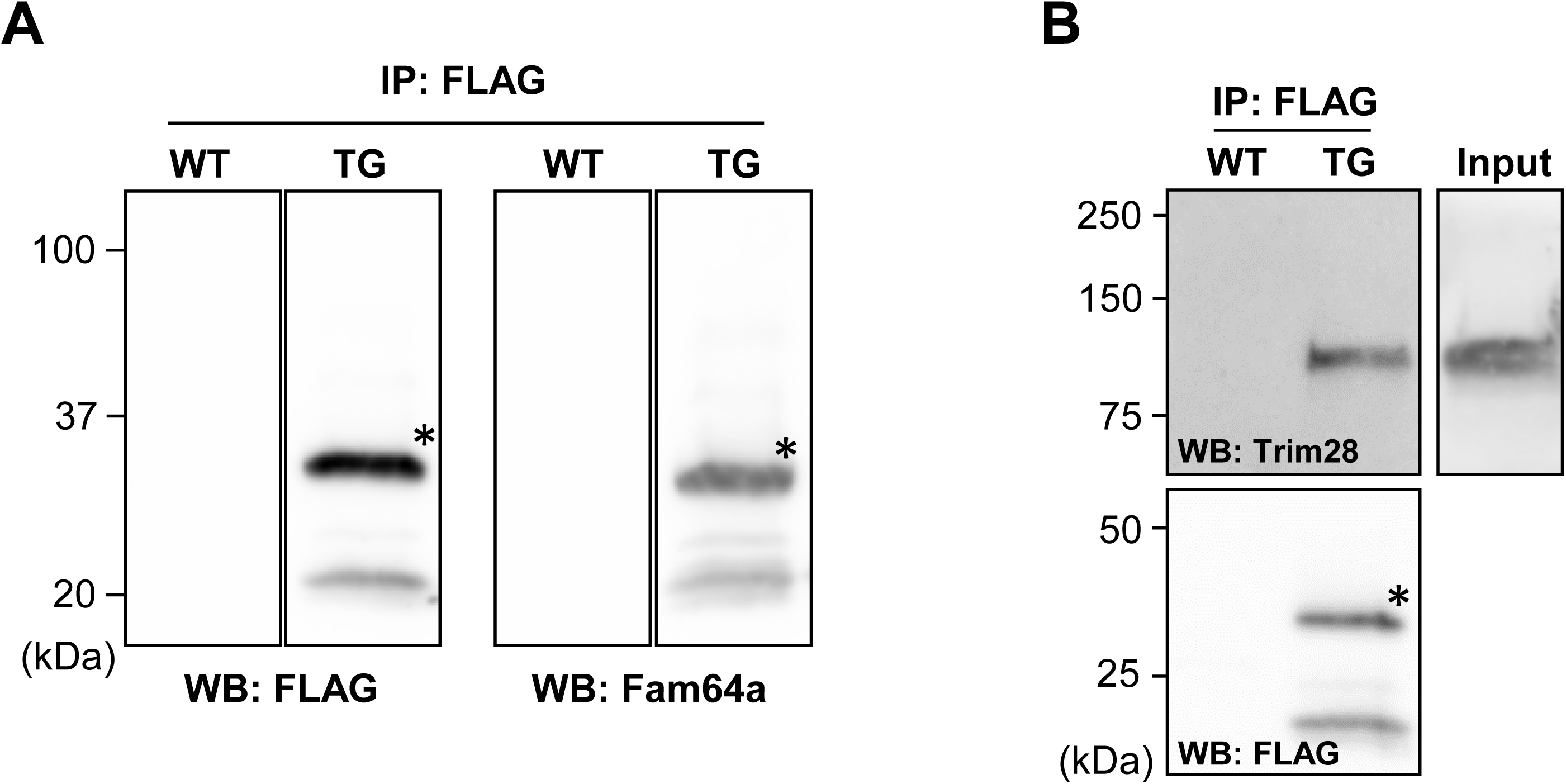
Comprehensive search for interacting partners of Fam64a. **A:** Immunoprecipitation (IP) against FLAG peptide that was expressed as a C-terminal tag of overexpressing Fam64a protein in TG mice hearts, followed by western blotting (WB) using FLAG and Fam64a antibody, which correctly detected Fam64a-FLAG fusion protein (*) in TG, but not in WT mice heart lysates, validating the immunoprecipitation procedure. **B:** The same IP/WB procedure was applied for Trim28, indicating that Fam64a forms complex with Trim28 in CMs.

## Notes

### Competing Interest Statement

The authors have declared no competing interest.

## References

1 Archangelo LF, Gläsner J, Krause A, Bohlander SK. The novel CALM interactor CATS influences the subcellular localization of the leukemogenic fusion protein CALM/AF10. Oncogene. 2006 Jul 6;25(29):4099–109. doi: 10.1038/sj.onc.1209438.

2 Archangelo LF, Greif PA, Maucuer A, Manceau V, Koneru N, Bigarella CL, Niemann F, dos Santos MT, Kobarg J, Bohlander SK, Saad ST. The CATS (FAM64A) protein is a substrate of the Kinase Interacting Stathmin (KIS). Biochim Biophys Acta. 2013 May;1833(5):1269–79. doi: 10.1016/j.bbamcr.2013.02.004.

3 Asada M, Rauch A, Shimizu H, Maruyama H, Miyaki S, Shibamori M, Kawasome H, Ishiyama H, Tuckermann J, Asahara H. DNA binding-dependent glucocorticoid receptor activity promotes adipogenesis via Krüppel-like factor 15 gene expression. Lab Invest. 2011 Feb;91(2):203–15. doi: 10.1038/labinvest.2010.170.

4 Barbutti I, Xavier-Ferrucio JM, Machado-Neto JA, Ricon L, Traina F, Bohlander SK, Saad ST, Archangelo LF. CATS (FAM64A) abnormal expression reduces clonogenicity of hematopoietic cells. Oncotarget. 2016 Oct 18;7(42):68385–68396. doi: 10.18632/oncotarget.11724.

5 Borden A, Kurian J, Nickoloff E, Yang Y, Troupes CD, Ibetti J, Lucchese AM, Gao E, Mohsin S, Koch WJ, Houser SR, Kishore R, Khan M. Transient introduction of miR-294 in the heart promotes cardiomyocyte cell cycle reentry after injury. Circ Res. 2019 Jun 21;125(1):14–25. doi: 10.1161/CIRCRESAHA.118.314223.

6 Chang CJ, Chen YL, Lee SC. Coactivator TIF1beta interacts with transcription factor C/EBPbeta and glucocorticoid receptor to induce alpha1-acid glycoprotein gene expression. Mol Cell Biol. 1998 Oct;18(10):5880–7. doi: 10.1128/mcb.18.10.5880.

7 Chiang CS, Huang CH, Chieng H, Chang YT, Chang D, Chen JJ, Chen YC, Chen YH, Shin HS, Campbell KP, Chen CC. The Ca(v)3.2 T-type Ca(2+) channel is required for pressure overload-induced cardiac hypertrophy in mice. Circ Res. 2009 Feb 27;104(4):522–30. doi: 10.1161/CIRCRESAHA.108.184051.

8 Cordeiro JM, Calloe K, Moise NS, Kornreich B, Giannandrea D, Di Diego JM, Olesen SP, Antzelevitch C. Physiological consequences of transient outward K+ current activation during heart failure in the canine left ventricle. J Mol Cell Cardiol. 2012 Jun;52(6):1291–8. doi: 10.1016/j.yjmcc.2012.03.001.

9 Cui M, Wang Z, Bassel-Duby R, Olson EN. Genetic and epigenetic regulation of cardiomyocytes in development, regeneration and disease. Development. 2018 Dec 20;145(24). pii: dev171983. doi: 10.1242/dev.171983.

10 Denslow SA, Wade PA. The human Mi-2/NuRD complex and gene regulation. Oncogene. 2007 Aug 13;26(37):5433–8. doi: 10.1038/sj.onc.1210611.

11 D’Uva G, Aharonov A, Lauriola M, Kain D, Yahalom-Ronen Y, Carvalho S, Weisinger K, Bassat E, Rajchman D, Yifa O, Lysenko M, Konfino T, Hegesh J, Brenner O, Neeman M, Yarden Y, Leor J, Sarig R, Harvey RP, Tzahor E. ERBB2 triggers mammalian heart regeneration by promoting cardiomyocyte dedifferentiation and proliferation. Nat Cell Biol. 2015 May;17(5):627–38. doi: 10.1038/ncb3149.

12 Fan L, Hsieh PN, Sweet DR, Jain MK. Krüppel-like factor 15: Regulator of BCAA metabolism and circadian protein rhythmicity. Pharmacol Res. 2018 Apr;130:123–126. doi: 10.1016/j.phrs.2017.12.018.

13 Fisch S, Gray S, Heymans S, Haldar SM, Wang B, Pfister O, Cui L, Kumar A, Lin Z, Sen-Banerjee S, Das H, Petersen CA, Mende U, Burleigh BA, Zhu Y, Pinto YM, Liao R, Jain MK. Kruppel-like factor 15 is a regulator of cardiomyocyte hypertrophy. Proc Natl Acad Sci U S A. 2007 Apr 24;104(17):7074–9. doi: 10.1073/pnas.0701981104.

14 Gabisonia K, Prosdocimo G, Aquaro GD, Carlucci L, Zentilin L, Secco I, Ali H, Braga L, Gorgodze N, Bernini F, Burchielli S, Collesi C, Zandonà L, Sinagra G, Piacenti M, Zacchigna S, Bussani R, Recchia FA, Giacca M. MicroRNA therapy stimulates uncontrolled cardiac repair after myocardial infarction in pigs. Nature. 2019 May;569(7756):418–422. doi: 10.1038/s41586-019-1191-6.

15 Han S, Zhang R, Jain R, Shi H, Zhang L, Zhou G, Sangwung P, Tugal D, Atkins GB, Prosdocimo DA, Lu Y, Han X, Tso P, Liao X, Epstein JA, Jain MK. Circadian control of bile acid synthesis by a KLF15-Fgf15 axis. Nat Commun. 2015 Jun 4;6:7231. doi: 10.1038/ncomms8231.

16 Hashimoto K, Kodama A, Honda T, Hanashima A, Ujihara Y, Murayama T, Nishimatsu SI, Mohri S. Fam64a is a novel cell cycle promoter of hypoxic fetal cardiomyocytes in mice. Sci Rep. 2017 Jun 30;7(1):4486. doi: 10.1038/s41598-017-04823-1.

17 Hashimoto K, Kodama A, Sugino M, Yobimoto T, Honda T, Hanashima A, Ujihara Y, Mohri S. Nuclear connectin novex-3 promotes proliferation of hypoxic foetal cardiomyocytes. Sci Rep. 2018 Aug 17;8(1):12337. doi: 10.1038/s41598-018-30886-9.

18 Honda T, Ujihara Y, Hanashima A, Hashimoto K, Tanemoto K, Mohri S. Turtle spongious ventricles exhibit more compliant diastolic property and possess larger elastic regions of connectin in comparison to rat compact left ventricles. Kawasaki Medical Journal 2018 44(1):1–17. doi: 10.11482/KMJ-E44(1)1

19 Ikeda S, Mizushima W, Sciarretta S, Abdellatif M, Zhai P, Mukai R, Fefelova N, Oka SI, Nakamura M, Del Re DP, Farrance I, Park JY, Tian B, Xie LH, Kumar M, Hsu CP, Sadayappan S, Shimokawa H, Lim DS, Sadoshima J. Hippo deficiency leads to cardiac dysfunction accompanied by cardiomyocyte dedifferentiation during pressure overload. Circ Res. 2019 Jan 18;124(2):292–305. doi: 10.1161/CIRCRESAHA.118.314048.

20 Jeyaraj D, Haldar SM, Wan X, McCauley MD, Ripperger JA, Hu K, Lu Y, Eapen BL, Sharma N, Ficker E, Cutler MJ, Gulick J, Sanbe A, Robbins J, Demolombe S, Kondratov RV, Shea SA, Albrecht U, Wehrens XH, Rosenbaum DS, Jain MK. Circadian rhythms govern cardiac repolarization and arrhythmogenesis. Nature. 2012 (a) Feb 22;483(7387):96–9. doi: 10.1038/nature10852.

21 Jeyaraj D, Scheer FA, Ripperger JA, Haldar SM, Lu Y, Prosdocimo DA, Eapen SJ, Eapen BL, Cui Y, Mahabeleshwar GH, Lee HG, Smith MA, Casadesus G, Mintz EM, Sun H, Wang Y, Ramsey KM, Bass J, Shea SA, Albrecht U, Jain MK. Klf15 orchestrates circadian nitrogen homeostasis. Cell Metab. 2012 (b) Mar 7;15(3):311–23. doi: 10.1016/j.cmet.2012.01.020.

22 Jiang L, Ren L, Zhang X, Chen H, Chen X, Lin C, Wang L, Hou N, Pan J, Zhou Z, Huang H, Huang D, Yang J, Liang Y, Li J. Overexpression of PIMREG promotes breast cancer aggressiveness via constitutive activation of NF-κB signaling. EBioMedicine. 2019;43:188–200. doi: 10.1016/j.ebiom.2019.04.001.

23 Jopling C, Sleep E, Raya M, Martí M, Raya A, Izpisúa Belmonte JC. Zebrafish heart regeneration occurs by cardiomyocyte dedifferentiation and proliferation. Nature. 2010 Mar 25;464(7288):606–9. doi: 10.1038/nature08899.

24 Karbassi E, Fenix A, Marchiano S, Muraoka N, Nakamura K, Yang X, Murry CE. Cardiomyocyte maturation: advances in knowledge and implications for regenerative medicine. Nat Rev Cardiol. 2020 Jun;17(6):341–359. doi: 10.1038/s41569-019-0331-x.

25 Katanosaka Y, Iwasaki K, Ujihara Y, Takatsu S, Nishitsuji K, Kanagawa M, Sudo A, Toda T, Katanosaka K, Mohri S, Naruse K. TRPV2 is critical for the maintenance of cardiac structure and function in mice. Nat Commun. 2014 May 29;5:3932. doi: 10.1038/ncomms4932.

26 Kaur K, Zangi L. Modified mRNA as a Therapeutic Tool for the Heart. Cardiovasc Drugs Ther. 2020 Dec;34(6):871–880. doi: 10.1007/s10557-020-07051-4.

27 Kita-Matsuo H, Barcova M, Prigozhina N, Salomonis N, Wei K, Jacot JG, Nelson B, Spiering S, Haverslag R, Kim C, Talantova M, Bajpai R, Calzolari D, Terskikh A, McCulloch AD, Price JH, Conklin BR, Chen HS, Mercola M. Lentiviral vectors and protocols for creation of stable hESC lines for fluorescent tracking and drug resistance selection of cardiomyocytes. PLoS One. 2009;4(4):e5046. doi: 10.1371/journal.pone.0005046.

28 Kubin T, Pöling J, Kostin S, Gajawada P, Hein S, Rees W, Wietelmann A, Tanaka M, Lörchner H, Schimanski S, Szibor M, Warnecke H, Braun T. Oncostatin M is a major mediator of cardiomyocyte dedifferentiation and remodeling. Cell Stem Cell. 2011 Nov 4;9(5):420–32. doi: 10.1016/j.stem.2011.08.013.

29 Kuo HC, Cheng CF, Clark RB, Lin JJ, Lin JL, Hoshijima M, Nguyêñ-Trân VT, Gu Y, Ikeda Y, Chu PH, Ross J, Giles WR, Chien KR. A defect in the Kv channel-interacting protein 2 (KChIP2) gene leads to a complete loss of I(to) and confers susceptibility to ventricular tachycardia. Cell. 2001 Dec 14;107(6):801–13. doi: 10.1016/s0092-8674(01)00588-8.

30 Kurokawa K, Shibasaki M, Mizuno K, Ohkuma S. Gabapentin blocks methamphetamine-induced sensitization and conditioned place preference via inhibition of α₂/δ-1 subunits of the voltage-gated calcium channels. Neuroscience. 2011 Mar 10;176:328–35. doi: 10.1016/j.neuroscience.2010.11.062.

31 Lee DS, Choi H, Han BS, Kim WK, Lee SC, Oh KJ, Bae KH. c-Jun regulates adipocyte differentiation via the KLF15-mediated mode. Biochem Biophys Res Commun. 2016 Jan 15;469(3):552–8. doi: 10.1016/j.bbrc.2015.12.035.

32 Leenders JJ, Wijnen WJ, Hiller M, van der Made I, Lentink V, van Leeuwen RE, Herias V, Pokharel S, Heymans S, de Windt LJ, Høydal MA, Pinto YM, Creemers EE. Regulation of cardiac gene expression by KLF15, a repressor of myocardin activity. J Biol Chem. 2010 Aug 27;285(35):27449–56. doi: 10.1074/jbc.M110.107292.

33 Leenders JJ, Wijnen WJ, van der Made I, Hiller M, Swinnen M, Vandendriessche T, Chuah M, Pinto YM, Creemers EE. Repression of cardiac hypertrophy by KLF15: underlying mechanisms and therapeutic implications. PLoS One. 2012;7(5):e36754. doi: 10.1371/journal.pone.0036754.

34 Locatelli P, Belaich MN, López AE, Olea FD, Uranga Vega M, Giménez CS, Simonin A, Bauzá MDR, Castillo MG, Cuniberti LA, Crottogini A, Cerrudo CS, Ghiringhelli PD. Novel insights into cardiac regeneration based on differential fetal and adult ovine heart transcriptomic analysis. Am J Physiol Heart Circ Physiol. 2020 Apr 1;318(4):H994–H1007. doi: 10.1152/ajpheart.00610.2019.

35 Mallipattu SK, Liu R, Zheng F, Narla G, Ma’ayan A, Dikman S, Jain MK, Saleem M, D’Agati V, Klotman P, Chuang PY, He JC. Kruppel-like factor 15 (KLF15) is a key regulator of podocyte differentiation. J Biol Chem. 2012 Jun 1;287(23):19122–35. doi: 10.1074/jbc.M112.345983.

36 Marshall LN, Vivien CJ, Girardot F, Péricard L, Scerbo P, Palmier K, Demeneix BA, Coen L. Stage-dependent cardiac regeneration in Xenopus is regulated by thyroid hormone availability. Proc Natl Acad Sci U S A. 2019 Feb 26;116(9):3614–3623. doi: 10.1073/pnas.1803794116.

37 Mohamed TMA, Ang YS, Radzinsky E, Zhou P, Huang Y, Elfenbein A, Foley A, Magnitsky S, Srivastava D. Regulation of cell cycle to stimulate adult cardiomyocyte proliferation and cardiac regeneration. Cell. 2018 Mar 22;173(1):104–116.e12. doi: 10.1016/j.cell.2018.02.014.

38 Nakada Y, Canseco DC, Thet S, Abdisalaam S, Asaithamby A, Santos CX, Shah AM, Zhang H, Faber JE, Kinter MT, Szweda LI, Xing C, Hu Z, Deberardinis RJ, Schiattarella G, Hill JA, Oz O, Lu Z, Zhang CC, Kimura W, Sadek HA. Hypoxia induces heart regeneration in adult mice. Nature. 2017 Jan 12;541(7636):222–227. doi: 10.1038/nature20173.

39 Oya E, Nakagawa R, Yoshimura Y, Tanaka M, Nishibuchi G, Machida S, Shirai A, Ekwall K, Kurumizaka H, Tagami H, Nakayama JI. H3K14 ubiquitylation promotes H3K9 methylation for heterochromatin assembly. EMBO Rep. 2019 Oct 4;20(10):e48111. doi: 10.15252/embr.201948111.

40 Peng W, Li M, Li H, Tang K, Zhuang J, Zhang J, Xiao J, Jiang H, Li D, Yu Y, Sham PC, Nattel S, Xu Y. Dysfunction of Myosin Light-Chain 4 (MYL4) Leads to heritable atrial cardiomyopathy with electrical, contractile, and structural components: evidence from genetically-engineered rats. J Am Heart Assoc. 2017 Oct 28;6(11). pii: e007030. doi: 10.1161/JAHA.117.007030.

41 Ray S, Pollard JW. KLF15 negatively regulates estrogen-induced epithelial cell proliferation by inhibition of DNA replication licensing. Proc Natl Acad Sci U S A. 2012 May 22;109(21):E1334–43. doi: 10.1073/pnas.1118515109.

42 Sasse SK, Mailloux CM, Barczak AJ, Wang Q, Altonsy MO, Jain MK, Haldar SM, Gerber AN. The glucocorticoid receptor and KLF15 regulate gene expression dynamics and integrate signals through feed-forward circuitry. Mol Cell Biol. 2013 Jun;33(11):2104–15. doi: 10.1128/MCB.01474-12.

43 Schultz DC, Friedman JR, Rauscher FJ 3rd. Targeting histone deacetylase complexes via KRAB-zinc finger proteins: the PHD and bromodomains of KAP-1 form a cooperative unit that recruits a novel isoform of the Mi-2alpha subunit of NuRD. Genes Dev. 2001 Feb 15;15(4):428–43. doi: 10.1101/gad.869501.

44 Stockdale WT, Lemieux ME, Killen AC, Zhao J, Hu Z, Riepsaame J, Hamilton N, Kudoh T, Riley PR, van Aerle R, Yamamoto Y, Mommersteeg MTM. Heart Regeneration in the Mexican Cavefish. Cell Rep. 2018 Nov 20;25(8):1997–2007.e7.doi: 10.1016/j.celrep.2018.10.072.

45 Strungs EG, Ongstad EL, O’Quinn MP, Palatinus JA, Jourdan LJ, Gourdie RG. Cryoinjury models of the adult and neonatal mouse heart for studies of scarring and regeneration. Methods Mol Biol. 2013;1037:343–53. doi: 10.1007/978-1-62703-505-7_20.

46 Sturzu AC, Rajarajan K, Passer D, Plonowska K, Riley A, Tan TC, Sharma A, Xu AF, Engels MC, Feistritzer R, Li G, Selig MK, Geissler R, Robertson KD, Scherrer-Crosbie M, Domian IJ, Wu SM. Fetal Mammalian Heart Generates a Robust Compensatory Response to Cell Loss. Circulation. 2015 Jul 14;132(2):109–21. doi: 10.1161/CIRCULATIONAHA.114.011490.

47 Taegtmeyer H, Sen S, Vela D. Return to the fetal gene program: a suggested metabolic link to gene expression in the heart. Ann N Y Acad Sci. 2010 Feb;1188:191–8. doi: 10.1111/j.1749-632.2009.05100.x.

48 Tashiro M, Hosokawa Y, Amao H, Tohei A. Duration of thermal support for preventing hypothermia induced by anesthesia with medetomidine-midazolam-butorphanol in mice. J Vet Med Sci. 2020 Dec 26;82(12):1757–1762. doi: 10.1292/jvms.20-0256.

49 Thomsen MB, Wang C, Ozgen N, Wang HG, Rosen MR, Pitt GS. Accessory subunit KChIP2 modulates the cardiac L-type calcium current. Circ Res. 2009 Jun 19;104(12):1382–9. doi: 10.1161/CIRCRESAHA.109.196972.

50 Ujihara Y, Iwasaki K, Takatsu S, Hashimoto K, Naruse K, Mohri S, Katanosaka Y. Induced NCX1 overexpression attenuates pressure overload-induced pathological cardiac remodelling. Cardiovasc Res. 2016 Sep;111(4):348–61. doi: 10.1093/cvr/cvw113.

51 Ujihara Y, Kanagawa M, Mohri S, Takatsu S, Kobayashi K, Toda T, Naruse K, Katanosaka Y. Elimination of fukutin reveals cellular and molecular pathomechanisms in muscular dystrophy-associated heart failure. Nat Commun. 2019 Dec 17;10(1):5754. doi: 10.1038/s41467-019-13623-2.

52 Umemura Y, Koike N, Ohashi M, Tsuchiya Y, Meng QJ, Minami Y, Hara M, Hisatomi M, Yagita K. Involvement of posttranscriptional regulation of Clock in the emergence of circadian clock oscillation during mouse development. Proc Natl Acad Sci U S A. 2017 Sep 5;114(36):E7479–E7488. doi: 10.1073/pnas.1703170114.

53 Wang WE, Li L, Xia X, Fu W, Liao Q, Lan C, Yang D, Chen H, Yue R, Zeng C, Zhou L, Zhou B, Duan DD, Chen X, Houser SR, Zeng C. Dedifferentiation, proliferation, and redifferentiation of adult mammalian cardiomyocytes after ischemic injury. Circulation. 2017 Aug 29;136(9):834–848. doi: 10.1161/CIRCULATIONAHA.116.024307.

54 Wei W, Lv Y, Gan Z, Zhang Y, Han X, Xu Z. Identification of key genes involved in the metastasis of clear cell renal cell carcinoma. Oncol Lett. 2019 May;17(5):4321–4328. doi: 10.3892/ol.2019.10130.

55 Yagita K, Horie K, Koinuma S, Nakamura W, Yamanaka I, Urasaki A, Shigeyoshi Y, Kawakami K, Shimada S, Takeda J, Uchiyama Y. Development of the circadian oscillator during differentiation of mouse embryonic stem cells in vitro. Proc Natl Acad Sci U S A. 2010 Feb 23;107(8):3846–51. doi: 10.1073/pnas.0913256107

56 Yao Z, Zheng X, Lu S, He Z, Miao Y, Huang H, Chu X, Cai C, Zou F. Knockdown of FAM64A suppresses proliferation and migration of breast cancer cells. Breast Cancer. 2019 Nov;26(6):835–845. doi: 10.1007/s12282-019-00991-2.

57 Yoda T, McNamara KM, Miki Y, Onodera Y, Takagi K, Nakamura Y, Ishida T, Suzuki T, Ohuchi N, Sasano H. KLF15 in breast cancer: a novel tumor suppressor? Cell Oncol (Dordr). 2015 Jun;38(3):227–35. doi: 10.1007/s13402-015-0226-8.

58 Zangi L, Oliveira MS, Ye LY, Ma Q, Sultana N, Hadas Y, Chepurko E, Später D, Zhou B, Chew WL, Ebina W, Abrial M, Wang QD, Pu WT, Chien KR. Insulin-Like Growth Factor 1 Receptor-Dependent Pathway Drives Epicardial Adipose Tissue Formation After Myocardial Injury. Circulation. 2017 Jan 3;135(1):59–72. doi: 10.1161/CIRCULATIONAHA.116.022064.

59 Zhang L, Prosdocimo DA, Bai X, Fu C, Zhang R, Campbell F, Liao X, Coller J, Jain MK. KLF15 establishes the landscape of diurnal expression in the heart. Cell Rep. 2015 Dec 22;13(11):2368–2375. doi: 10.1016/j.celrep.2015.11.038.

60 Zhao WM, Coppinger JA, Seki A, Cheng XL, Yates JR 3rd, Fang G. RCS1, a substrate of APC/C, controls the metaphase to anaphase transition. Proc Natl Acad Sci U S A. 2008 Sep 9;105(36):13415–20. doi: 10.1073/pnas.0709227105.

61 Zhu Y, Do VD, Richards AM, Foo R. What we know about cardiomyocyte dedifferentiation. J Mol Cell Cardiol. 2020 Dec 1;152:80–91. doi: 10.1016/j.yjmcc.2020.11.016.

